# A single extracellular loop of FoxA controls ligand specificity, uptake, and signaling in *Pseudomonas aeruginosa*

**DOI:** 10.1101/2022.11.18.517105

**Authors:** Derek C. K. Chan, Inokentijs Josts, Kalinka Koteva, Gerard D. Wright, Henning Tidow, Lori L. Burrows

## Abstract

The outer membrane of Gram-negative bacteria prevents many antibiotics from reaching intracellular targets. However, some antimicrobials can take advantage of iron import transporters to cross this barrier. We showed previously that the thiopeptide antibiotic, thiocillin, exploits the nocardamine (ferrioxamine E) xenosiderophore transporter, FoxA, of the opportunistic pathogen *Pseudomonas aeruginosa* for uptake. Here we show that FoxA also transports the xenosiderophore bisucaberin and describe at 2.5 Å resolution the first crystal structure of bisucaberin bound to FoxA. Bisucaberin is distinct from other siderophores because it forms a 3:2 rather than 1:1 siderophore-iron complex. Mutations in a single extracellular loop of FoxA differentially affected nocardamine, thiocillin, and bisucaberin binding, uptake, and signal transduction. These results show that in addition to modulating ligand binding, the extracellular loops of siderophore transporters are of fundamental importance for controlling ligand uptake and its regulatory consequences, which has implications for the development of siderophore-antibiotic conjugates to treat difficult infections.

---

Iron is an essential micronutrient for bacteria, involved in biofilm formation^1^, virulence factor production^2^, and colonization and infection^3^. Bacteria have evolved ways to survive in iron-limited conditions such as those encountered at sites of infection^4,5^. One common strategy includes the production and release of siderophores into the environment. Siderophores are natural products with high affinity for iron, and ferri-siderophore complexes are taken up via specific transporters in the outer membrane (OM)^6–9^. The opportunistic human pathogen, *Pseudomonas aeruginosa*, makes two main siderophores, pyoverdine and pyochelin^10–12^ and mutants deficient in production of those siderophores are unable to grow in iron-limited conditions^6^. Transport of siderophores into the cell is an energy-dependent process^7^ that relies on the inner-membrane TonB-ExbD-ExbD complex (Fig. 1A) to couple the proton motive force (PMF) to TonB-dependent transporters (TBDTs)^14,15^. TBDTs are 22-stranded β-barrel proteins occluded by a central plug domain that prevents nonspecific diffusion. *P. aeruginosa* encodes dozens of predicted TBDTs that recognize native siderophores and xenosiderophores produced by other microorganisms.

**Fig. 1.**
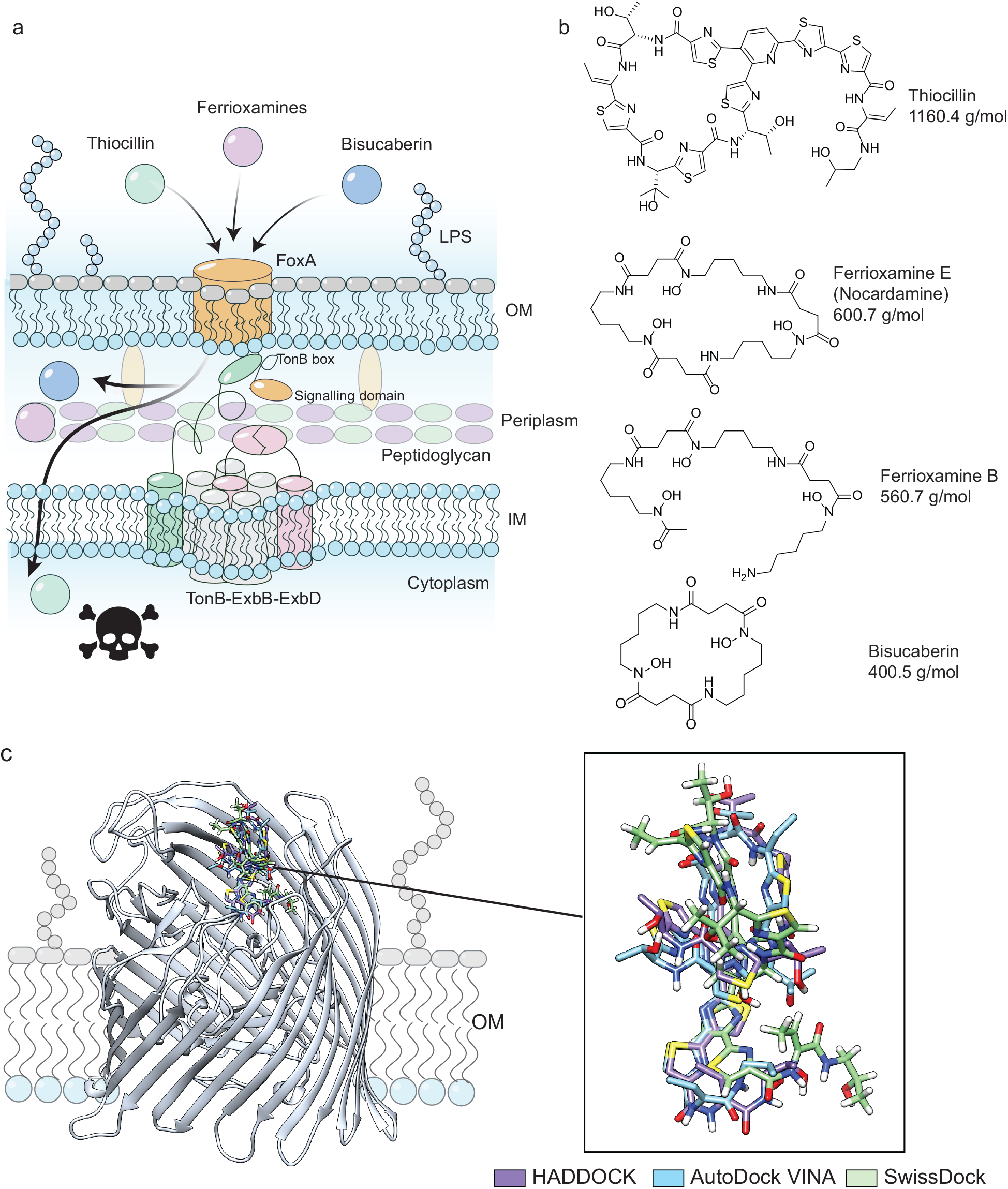
Thiocillin hijacks FoxA to cross the outer membrane. **a,** Model of thiocillin and siderophore uptake. OM = outer membrane. IM = inner membrane. LPS = lipopolysaccharide. **b,** Structures of thiocillin, nocardamine, ferrioxamine B, and bisucaberin. Siderophores are shown in their apo state. **c,** Superimposed docking predictions of thiocillin with apo-FoxA (PDB 6I97). Stick models of thiocillin docked into FoxA (ribbon) using different docking algorithms are shown in purple (HADDOCK), blue (AutoDock VINA), and green (SwissDock). All three docking simulations converge at a similar binding site. Heteroatoms are depicted as different colours.

Extracellular loops connecting the β-strands of TBDTs are involved in ferri-siderophore binding^7^. Upon ligand binding, the transporter undergoes conformational changes, including the inward movement of an extracellular loop^13,16^ and exposure of a TonB-box motif on the periplasmic face of the plug that interacts with TonB^13^. Some transporters also have an extended periplasmic N-terminus containing a signaling domain^17–19^. In a feed-forward loop, ligand uptake via the transporter triggers the release of a σ/anti-σ pair that recruits RNA polymerase to the promoter of the relevant TBDT^20^ to promote its increased expression.

Some bacteriophages^21^, bacteriocins^22,23^, lassopeptides^24,25^, and antibiotics^26,27^ use TBDTs to cross the OM, although the details of uptake remain unclear. The ability of TBDT-dependent antimicrobials to bypass the OM barrier, coupled with their narrow-spectrum activity, makes them interesting candidates for human use. Previously, we showed that the thiopeptide antibiotic thiocillin requires the FoxA transporter to inhibit growth of *P. aeruginosa*^28^ (Fig. 1a). Thiocillin is a cyclic thiazole-containing antibiotic (Fig. 1b) that binds the ribosome at the interface of the L11 protein and elongation factor G to inhibit protein synthesis^29^. FoxA is the TBDT for the siderophores ferrioxamine B and nocardamine (ferrioxamine E)^6,13,20^ (Fig. 1b), which prior to our work were its only known ligands. This work focuses on nocardamine as it exclusively uses FoxA for uptake^6^ whereas ferrioxamine B uses both FoxA and FpvB^30^. *P. aeruginosa* FoxA has an extended N-terminus containing a signaling domain^19,20^ involved in upregulating FoxA expression in response to ligand binding through release of anti-σ/σ factor pair FoxI/FoxR^18,20^. However, the way in which thiocillin interacts with FoxA and whether it triggers conformational changes and signaling events similar to those induced by nocardamine are unknown.

Here we show that binding, uptake, and signaling through FoxA is dependent on the nature of the ligand and controlled by a mobile extracellular loop. Besides thiocillin and nocardamine, FoxA also acts as a low-affinity transporter for the nocardamine-like hydroxamate siderophore, bisucaberin (Fig. 1b). The co-crystal structure of a 3:2 bisucaberin-iron complex with FoxA was determined at 2.5 Å resolution^31^, showing that bisucaberin mimics the binding conformation of nocardamine but interacts with unique residues, similar to thiocillin. Despite these differences, we show that extracellular loop 8 (L8) of FoxA modulates uptake and signaling for all three ligands. The conservation of this loop configuration among TBDTs suggests that it may play a similar role in other transporters.

## RESULTS

### At least five residues in FoxA are essential for thiocillin susceptibility

After unsuccessful co-crystallization attempts, we took a molecular docking approach to probe the potential interactions between thiocillin and FoxA. Docking simulations using AutoDock VINA^32^, SwissDock^33^, and HADDOCK^34^ were compared (Fig. 1c). All three algorithms predicted that thiocillin binds in the same region of FoxA as nocardamine, with AutoDock VINA and HADDOCK predicting similar poses. Therefore, subsequent experiments to verify predicted interactions by site-directed mutagenesis were based on those results (Fig. 2a). Using a low copy number arabinose-inducible expression vector, pHERD20T^35^, 83 single residue mutant FoxA transporters were generated based on the predictions, expressed in a *P. aeruginosa* PA14 Δ*foxA* background, and screened for a) restoration of thiocillin susceptibility in 10:90 medium, and b) the ability to grow in iron-limited casamino acids (CAA) medium supplemented with nocardamine. Mutants unable to make FoxA are resistant to thiocillin^28^ and growth is inhibited by nocardamine through iron restriction since they cannot take up the siderophore. Complementation with wild-type (WT) FoxA restores thiocillin susceptibility and growth with nocardamine.

**Fig. 2.**
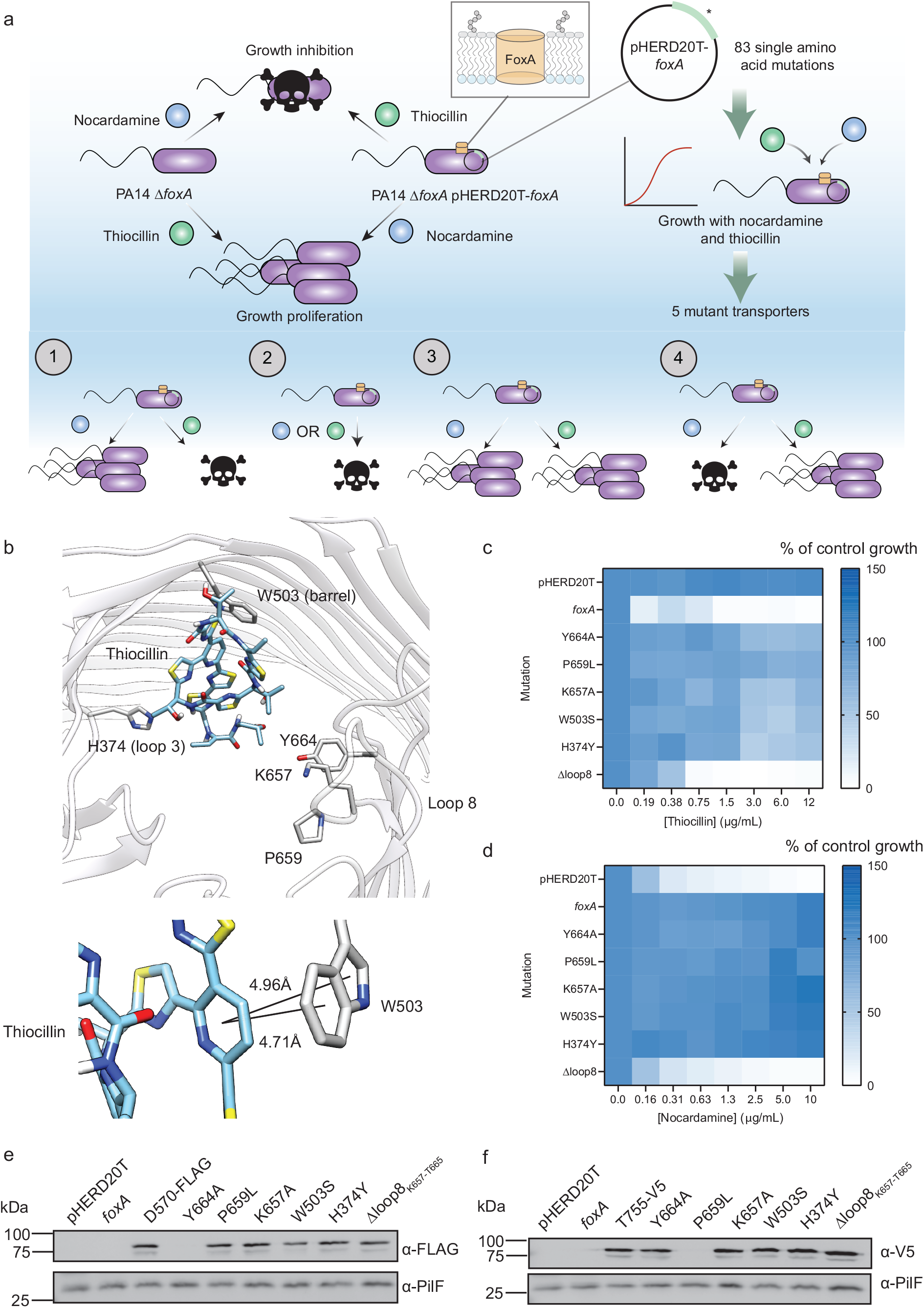
Identifying FoxA residues important for thiocillin activity and nocardamine uptake. **a,** Schematic depicting dual growth phenotype screening for the identification of single residues important for nocardamine and thiocillin uptake through FoxA. In iron-limited media, *P. aeruginosa* Δ*foxA* is susceptible to growth inhibition by nocardamine but resistant to thiocillin. Complementation of the mutant *in trans* with a WT copy of *foxA* restores growth with nocardamine and susceptibility to thiocillin. Based on the docking simulations, 83 single amino acid mutations hypothesized to be important for ligand interaction were made and expressed *in trans* for screening. Of these mutant transporters, five failed to restore thiocillin susceptibility but supported growth with nocardamine. From the screen, four categories of mutant were observed. Category 1: the transporter permits uptake of both ligands similar to WT. Category 2: the transporter restores thiocillin susceptibility but does not support growth with nocardamine. Category 3: the transporter does not restore thiocillin susceptibility but supports growth with nocardamine. Category 4: the transporter is either not expressed or non-functional. **b,** (Top) Stick model of thiocillin docked into FoxA (ribbon model) with residues identified as important for thiocillin activity shown. (Bottom) A zoomed-in view of the predicted interaction between the central pyridine ring of thiocillin and FoxA W503 with distances shown. **c,** Thiocillin MIC assays with Δ*foxA* expressing WT or mutant FoxA transporters and **d,** nocardamine growth assays. A darker shade of blue indicates more growth and a lighter shade, less growth. White indicates no growth. Growth is expressed as percent of control of the vehicle control (DMSO). Results are an average of three biological replicates. **e, f,** Blots of WT and mutant FoxA expression *in trans*. Empty vector and unlabeled WT-FoxA were included as negative controls. Internal FLAG (D570FLAG) and V5 (T755V5) tagged WT and mutant FoxA receptors are expressed at similar levels. PilF was used as a loading control for outer membrane proteins.

Y664A, P659L, K657A (L8), W503S (barrel), and H374Y (loop 3) mutants failed to restore thiocillin susceptibility (Fig. 2b-c) but allowed growth with the siderophore (Fig. 2d). The data were consistent with the docking predictions, where W503 interacts with the pyridine ring of thiocillin through π-stacking (Fig. 2b). Unexpectedly, truncation of L8_K657-T665_ was permissive for thiocillin uptake, but prevented growth with nocardamine (Fig. 2c,d). To monitor stability of the mutant transporters, WT FoxA was tagged with V5 or FLAG epitope tags at one of 32 different locations to identify functionally permissive sites, and each variant tested for its ability to restore WT levels of thiocillin susceptibility and growth with nocardamine in the Δ*foxA* background (Extended Data Fig. 1a,b). Then, each point mutant transporter was similarly tagged to monitor expression levels. T755-V5 and D570-FLAG variants were selected for monitoring, because the tags were located on loops distinct from those containing residues important for thiocillin uptake. Outer-membrane fractions were isolated and the levels of WT and mutant transporters assessed by Western blot (Fig. 2e,f). PilF, an outer membrane lipoprotein necessary for type IV pilus secretin localization and multimerization^36^ was used as a loading control. Although Y664A T755-V5, and P659L D570-FLAG could not be detected, the alternately tagged Y664A D570-FLAG and P659L T755-V5 were detectable, confirming that they are expressed. All other receptors were expressed at levels similar to the tagged WT.

### Nocardamine and thiocillin compete for binding and uptake

Thiocillin is predicted to bind in the same pocket as nocardamine (Fig. 3a,b), suggesting that two ligands could compete for the transporter. Consistent with this prediction, antagonism was observed in checkerboard assays with thiocillin and nocardamine-Fe^3+^ against Δ*foxA* complemented with WT FoxA *in trans* (Fig. 3c). Nocardamine also antagonized thiocillin uptake, although >100-fold higher concentrations were required compared to the iron-bound form (Extended Data Fig. 2a). Thiocillin antagonism was observed only with nocardamine, not other siderophores, supporting the model of competition for similar binding sites (Extended Data Fig. 2b-d).

**Fig. 3.**
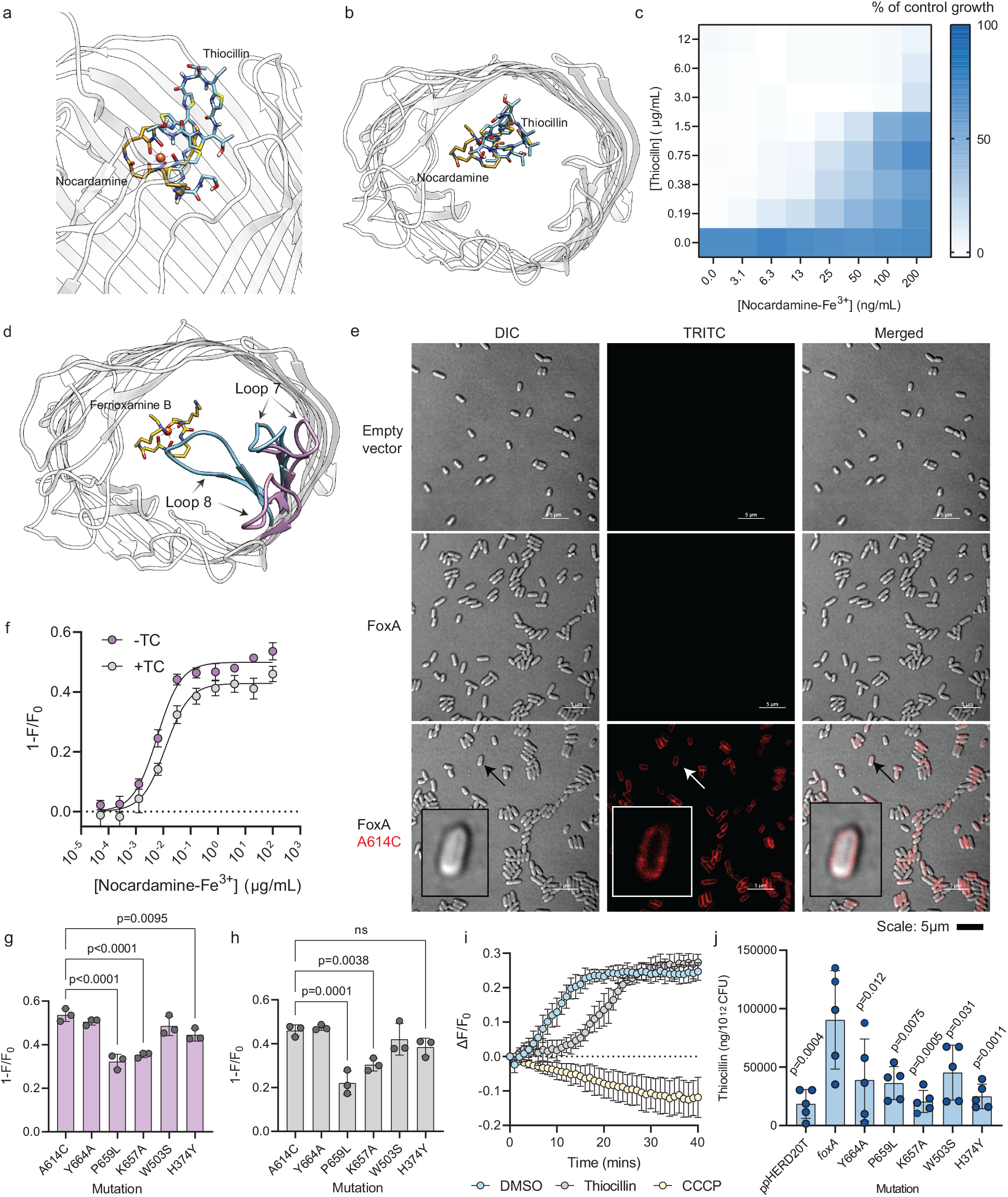
Thiocillin competes with nocardamine-Fe^3+^ for FoxA. **a,** Superimposed structures of nocardamine-Fe^3+^ (PDB:6Z8A) and thiocillin (AutoDock Vina) inside FoxA. Side view and **b,** top view are shown. Both compounds are shown as stick models, with nocardamine in yellow and thiocillin in blue. **c,** Checkerboard assay of PA14 Δ*foxA* pHERD20T-*foxA* treated with thiocillin and nocardamine-Fe^3+^. Results are averaged from three independent biological replicates. **d,** L7 and L8 movement upon ferrioxamine B-Fe^3+^ binding (PDB 6I96 and 6I97). The purple loops indicate the orientation prior to ferri-siderophore binding while blue loops indicate the orientation after binding. **e,** Fluorescence microscopy images of PA14 Δ*foxA* with pHERD20T, *foxA*, or *foxA* A614C labeled with AlexaFluor 594. Inset: zoomed-in view of a single labeled cell. Scale bar is 5μm. **f,** Fluorescence quenching of A614C-labeled FoxA after the addition of nocardamine-Fe^3+^ without thiocillin pre-treatment (purple) or with thiocillin pre-treatment (grey). Results are averaged from three independent biological replicates. K_d_ was calculated in GraphPad Prism with a one-site binding model. A614C is highlighted in red to indicate the labeled residue. **g,** Fluorescence quenching of A614C-labeled FoxA at 100 μg/mL nocardamine-Fe^3+^ in the absence or **h,** presence of thiocillin pre-treatment. Statistics were calculated with GraphPad Prism using one-way ANOVA followed by Dunnett’s multiple comparisons test; p values are shown, ns = not significant. Results from three independent biological replicates are shown. **i,** Fluorescence recovery of PA14 Δ*foxA* pHERD20T-*foxA* A614C labeled with fluorescein-5-maleimide treated with 6.4 ng/mL nocardamine-Fe^3+^ with DMSO (blue), 12 μg/mL (10 μM) thiocillin (grey), and 100 μM CCCP. Results are averaged from three independent biological replicates. **j,** Uptake of thiocillin through FoxA or its single amino acid mutants. Thiocillin concentrations are normalized to ng of thiocillin per 10^12^ CFU. Results were averaged from five independent biological replicates. Statistics were calculated with GraphPad Prism using one-way ANOVA followed by Dunnett’s multiple comparisons test; p-values are shown.

A FoxA whole-cell fluorescent sensor was developed to detect nocardamine-Fe^3+^ binding and uptake. Previous studies used site-specific cysteine labeling to incorporate fluorescein-5-maleimide at extracellular loops to detect ligand binding^37,38^. The fluorescence of the labeled TBDT can be quenched with increasing concentrations of siderophore-Fe^3+^ due to conformational changes that alter the environment surrounding the fluorophore. Comparison of apo- and ferrioxamine B-bound FoxA structures showed that loop 7 (L7) and L8 fold downwards toward the ligand upon binding (Fig. 3d, Extended Data Fig. 3a, Supplementary Data File 1)^13^; therefore, 19 amino acid residues in those loops were individually mutated to Cys and screened for those that could be labeled for detection (Extended Data Fig. 3a). The Cys mutant transporters were expressed in the Δ*foxA* background and screened for functionality based on their ability to complement thiocillin susceptibility and growth with nocardamine (Extended Fig. 3b,c). Of 17 functional Cys point mutants, we observed fluorescence quenching by nocardamine-Fe^3+^ for Q660C (L8) and A614C (L7) (Extended Data Fig. 3d). Fluorescence of labeled A614C and Q660C FoxA was significantly greater compared to WT FoxA and labeling did not affect thiocillin susceptibility (Extended Data Fig. 3e,f). A614C was selected for further experiments to avoid potential steric interactions with residues important for thiocillin uptake. Interestingly, no fluorescence quenching was observed for thiocillin or apo-nocardamine even at 12μg/mL (10μM) and 200μg/mL respectively, suggesting that the two ligands induce conformational changes different from those caused by nocardamine-Fe^3+^ (Extended Data Fig. 3g,h). A614C-labeling was also specific, as fluorescence was localized to the periphery of cells (Fig. 3e). A614C was introduced into the single residue FoxA mutants and expressed in Δ*foxA* to confirm that the Cys mutation had no effect on thiocillin susceptibility or growth with nocardamine compared to the single residue mutations alone (Extended Data Fig. 4a,b). In the absence of thiocillin, the K_d_ of nocardamine-Fe^3+^ was 6.0 ± 0.62 ng/mL (Fig. 3f, Extended Data Fig. 5). Since thiocillin failed to quench fluorescence, we could isolate its impact on the binding affinity of nocardamine-Fe^3+^. Pre-treating cells with 12μg/mL thiocillin for 5 min increased the K_d_ to 12 ± 1.8 ng/mL, indicating weak competition, consistent with the results of checkerboard assays that demonstrated antagonism between the two ligands. As a control, 11 μg/mL (10 μM) of geninthiocin A, a thiopeptide that lacks activity against *P. aeruginosa*, was tested and the K_d_ was similar to that of nocardamine-Fe^3+^ alone (Extended Data Fig. 4c).

The effects on K_d_ of the single residue mutations and the L8 deletion were also determined using A614C labeled transporters (Extended Data Fig. 4d-i, Extended Data Fig. 5, Extended Data Fig. 6). The K_d_ of nocardamine-Fe^3+^ for Y664A, P659L, K657A, and W503S was similar to the WT in the absence of thiocillin (Extended Data Fig. 4d-h, Extended Data Fig. 5); however, H374Y increased the K_d_ 26-fold. This increase was expected because H374 forms a H-bond with the hydroxamate side chain of nocardamine^6,13^. Changing that residue to tyrosine may displace the siderophore-iron complex in the binding pocket, reducing its interactions with other side chains. Thiocillin pre-treatment increased the K_d_ of nocardamine-Fe^3+^ for all the single residue mutants. P659L and K657A had 14- and 8-fold increases with thiocillin treatment, respectively, compared to the WT (Extended Data Fig. 5). Unexpectedly, P659L and K657A also showed significantly reduced quenching, possibly due to reduced loop movement since the K_d_s are similar to the WT (Fig. 3g,h, Extended Data Fig. 5). FoxA_ΔK657-T665_ could still bind nocardamine-Fe^3+^ with a 14-fold increase in K_d_ even though uptake of the ferri-siderophore was compromised (Fig. 2d, Extended Data Fig. 4i, Extended Data Fig. 5).

Thiocillin competed with nocardamine for uptake as well as binding. Fluorescence recovery was used as an indicator of uptake^38^ as the protein reverts to the apo-state once the siderophore is released into the periplasm. As a control, cells were also incubated with 100 μM carbonyl cyanide-m-chlorophenyl hydrazone (CCCP), a PMF uncoupler, to prevent uptake. The sensor strain treated with 6.4 ng/mL of nocardamine-Fe^3+^ recovered 50% of baseline fluorescence after 8.9 min (Fig. 3I). Cells pretreated with thiocillin took 20 min to recover 50% fluorescence, more than twice as long. CCCP treatment inhibited fluorescence recovery and continuous quenching was observed, suggesting that once bound, the ligand must be taken up to allow the transporter to return to its apo confirmation.

Since thiocillin increased the K_d_ for nocardamine in WT FoxA and the mutants, we inferred that the thiopeptide could bind the mutant transporters. This result suggested that the reduced susceptibility to thiocillin is due to reduced uptake rather than binding. A mass spectrometry approach was used to test this, measuring intracellular thiocillin accumulation after treating with 12 μg/mL thiocillin for 1 h. Thiocillin concentrations in Δ*foxA* complemented with WT FoxA were significantly greater than the vector control or when Δ*foxA* was complemented with the mutant transporters, supporting the hypothesis that the mutations reduce thiocillin uptake (Fig. 3j).

### Residues that impact thiocillin uptake are important for stimulation of FoxA expression by nocardamine

Since mutations that affected thiocillin uptake also reduced nocardamine-Fe^3+^ binding, they could potentially impact signaling (Fig. 4a). Binding of nocardamine to FoxA induces conformational changes that lead to the degradation and release of an anti-σ/σ factor pair (FoxI/FoxR). FoxR then guides RNA polymerase to the promoter of *foxA* to upregulate its transcription. The exact mechanism of signal transduction is unknown, but we hypothesized that the residues we identified may be important. We tested signalling using a promoter assay where GFP expression is under control of P*_foxA_*. Chromosomal knock-in mutants encoding FoxA Y664A, P659L, K657A, W503S, or H374Y were generated so the proteins were expressed from the native promoter. P*_foxA_-gfp* and a promoterless control (P_x_-*gfp*) were introduced into the WT and each mutant. The strains were treated with increasing concentrations of thiocillin (Fig. 4b,c) or nocardamine (Fig. 4d,e). Thiocillin did not induce GFP expression. When treated with nocardamine, W503S and Y664A had reduced GFP expression compared to the WT, while GFP production from P659L, K657A, and H374Y was abolished, similar to the promoterless negative control. GFP expression in all the mutants was significantly lower than WT at the highest concentration of nocardamine tested (Fig. 4f). These results suggest that the mutated residues are important in the signaling response. The difference in signaling between thiocillin and nocardamine further indicates that the protein conformations that result from binding of these ligands are different, consistent with the results of mutational analyses and fluorescence quenching assays.

**Fig. 4.**
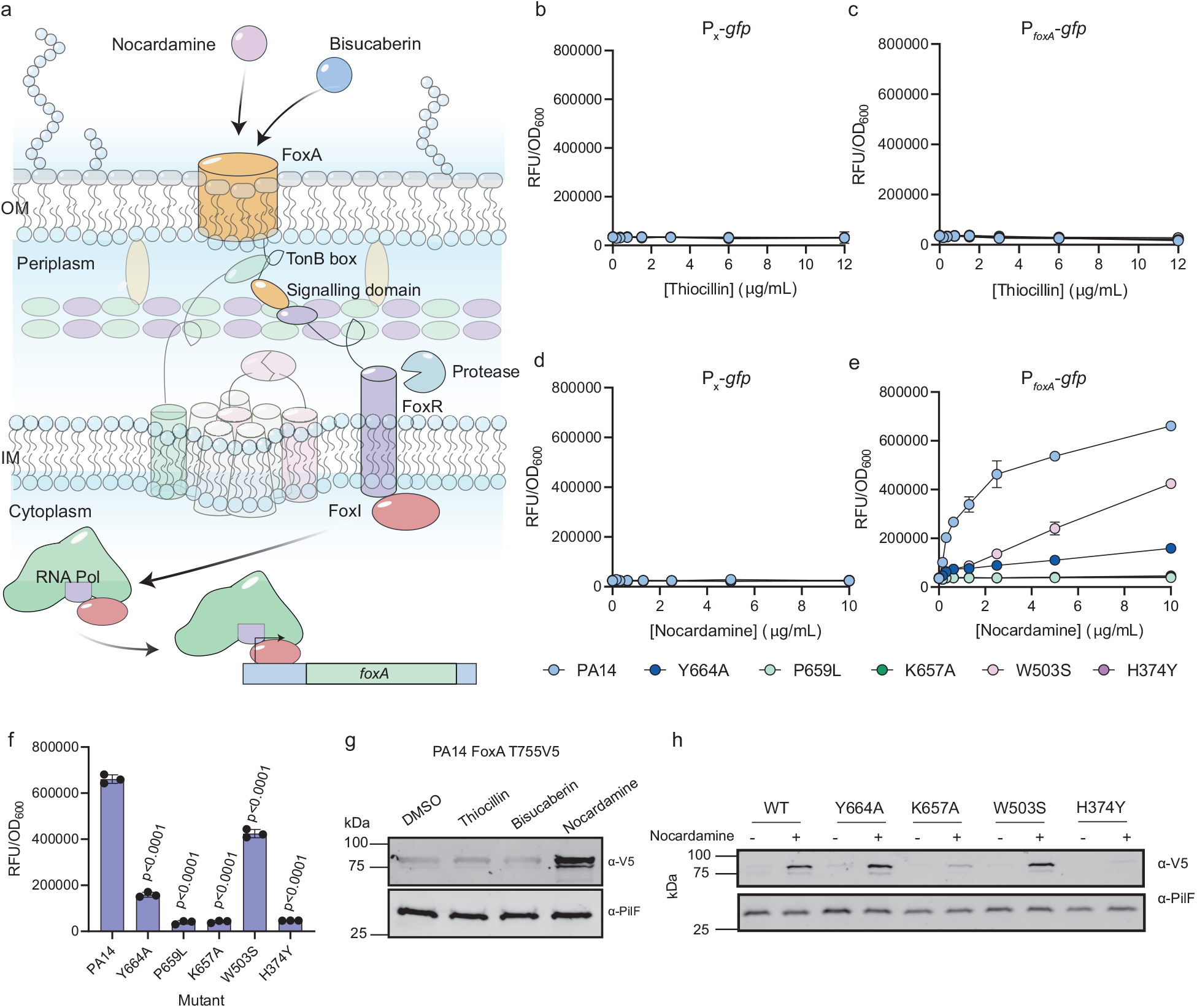
FoxA single residue mutants are defective in nocardamine signaling. **a,** Diagram depicting nocardamine-induced expression of FoxA by the FoxI/FoxR σ/anti-σ factors. Binding and uptake of nocardamine triggers conformational changes in FoxA, including exposure of the TonB box motif and signalling domain to the FoxR anti-sigma factor. Proteolytic cleavage of FoxR in the periplasm leads to the release of FoxI into the cytoplasm, which directs RNA polymerase to P*_foxA_*. **b,** WT PA14 and chromosomal FoxA mutants harbouring a promoterless GFP plasmid and **c,** P*_foxA_* treated with increasing concentrations of thiocillin in 10:90. Results are averaged from three independent biological replicates. WT PA14 and chromosomal FoxA mutants harbouring **d,** a promoterless GFP plasmid and **e,** P*_foxA_* treated with increasing concentrations of nocardamine in CAA. Results are averaged from three independent biological replicates. **f,** RFU/OD_600_ values for WT PA14 and FoxA chromosomal mutants treated with 10 μg/mL nocardamine in CAA. p values are shown. **g,** PA14 FoxA T775-V5 treated with 5 μg/mL thiocillin, bisucaberin, or nocardamine in CAA. Outer membranes were isolated and probed with an α-V5 antibody and α-PilF was used as a loading control. **h,** PA14 FoxA V5-tagged chromosomal single amino acid mutants were treated with 5 μg/mL nocardamine in CAA overnight. Outer membranes were purified and probed with an α-V5 antibody and α-PilF was used as a loading control.

We next looked at FoxA protein expression. FoxA T755-V5 was chromosomally integrated to allow for detection of expression by Western blot and the tagged strain treated with 5.0 μg/mL each of thiocillin, bisucaberin – a cyclic bis-hydroxamate siderophore structurally related to nocardamine but lacking a hydroxamate group (Fig. 1b) – or nocardamine (Fig. 4g). Only nocardamine stimulated FoxA expression (Fig. 4g). The tagged point mutants were also chromosomally integrated (except for P659L T775-V5 since it was not detectable when expressed *in trans*; Fig. 2f) and treated with 5.0 μg/mL nocardamine. WT, Y664A, and W503S variants had detectable expression, consistent with the promoter reporter assays (Fig. 4h). These data show there is overlap between residues important for thiocillin uptake and signaling by the siderophore (Extended Data Fig. 7).

### Bisucaberin exclusively uses the FoxA receptor for uptake

We initially tested bisucaberin because it was a hydroxamate siderophore with structural similarity to nocardamine. However, based on the expression induction assay, it did not appear to use FoxA for uptake in WT cells (Fig. 4g). Unlike the 1:1 nocardamine:iron complex, bisucaberin forms a 3:2 complex at physiological pH^31^ which could impact recognition of the complex by FoxA (Fig. 5a). To test this, we generated a *P. aeruginosa* PA14 mutant unable to make its native siderophores pyoverdine and pyochelin, Δ*pvdA* Δ*pchA*. This mutant requires exogenously-supplemented siderophores to grow in iron-limited media. Δ*pvdA* Δ*pchA* grew in CAA when provided with bisucaberin, suggesting that the siderophore was taken up (Fig. 5b). A previous study suggested that alcaligin, a siderophore similar to bisucaberin, may be taken up via the uncharacterized transporter PA14_46640^39^. To identify the transporter for bisucaberin, we deleted *PA14_46640* or *foxA* in the Δ*pvdA* Δ*pchA* background and grew the mutants with bisucaberin (Fig. 5b). The Δ*pvdA* Δ*pchA* Δ*PA14*_*46640* triple mutant grew similarly to WT when provided with bisucaberin; however, loss of *foxA* prevented growth, suggesting that bisucaberin uses FoxA exclusively for uptake in the absence of pyoverdine and pyochelin.

**Fig. 5.**
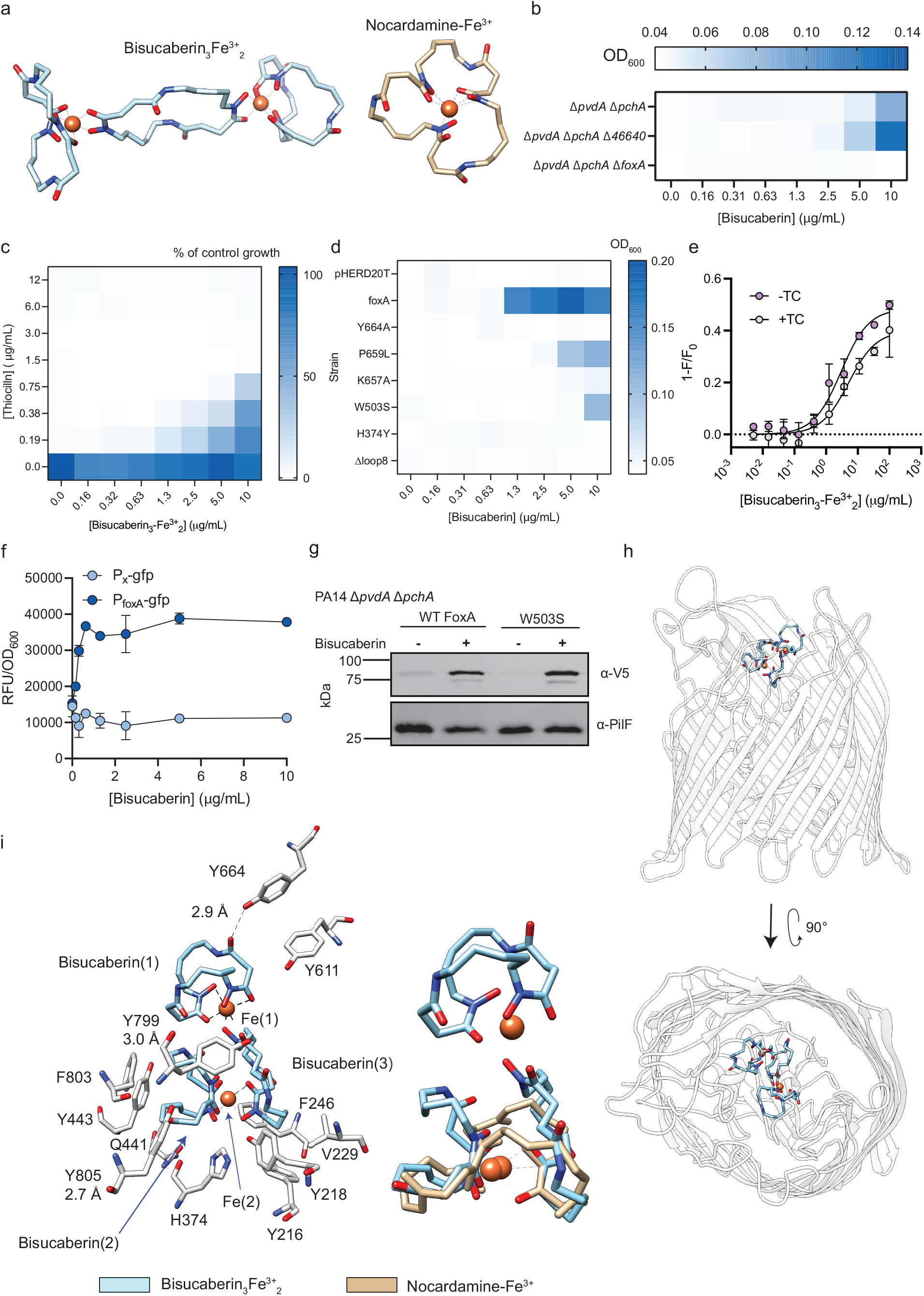
FoxA is a low affinity transporter for bisucaberin that induces signal transduction through FoxA. **a,** Structures of bisucaberin (left) and nocardamine (right) iron complexes modeled based on the mono-bridged complex of ferro-alcaligin and PDB: 6Z8A with FoxA removed respectively. Heteroatoms are in different colours: blue: nitrogen, red: oxygen, and Fe^3+^ is depicted as an orange sphere. **b,** Growth recovery of PA14 Δ*pvdA* Δ*pchA*, Δ*pvdA* Δ*pchA* Δ*PA14*_*46640*, and Δ*pvd A*Δ*pchA* Δ*foxA* treated with bisucaberin in CAA supplemented with increasing concentrations of bisucaberin. Legend shows OD_600_ where blue indicates growth and white indicates no growth. **c,** Checkerboard assay with PA14 Δ*pvdA* Δ*pchA* Δ*foxA* pHERD20T-*foxA* treated with thiocillin and bisucaberin-Fe^3+^ in 10:90 + 1% arabinose. **d,** Growth recovery of Δ*pvdA* Δ*pchA* Δ*foxA* overexpressing different FoxA mutant transporters treated with bisucaberin in CAA + 1% arabinose supplemented with increasing concentrations of bisucaberin. **e,** Fluorescence quenching of PA14 Δ*pvdA* Δ*pchA* Δ*foxA* pHERD20T-*foxA* A614C with bisucaberin-Fe^3+^ in the absence (purple) and presence (grey) of thiocillin. **f,** Promoter assays with Δ*pvdA* Δ*pchA* P*_x_-gfp* (light blue) and P*_foxA_-gfp* (dark blue) treated with bisucaberin in 10:90 medium supplemented with bisucaberin. Raw fluorescence values are normalized to OD_600_. Growth, fluorescence quenching, and promoter assays were averaged from three independent biological replicates. **g,** FoxA expression of chromosomally tagged FoxA and FoxA mutants treated with DMSO (−) or with 5.0 μg/mL bisucaberin (+). PilF was used as the loading control. A representative blot is shown. **h,** Structure of bisucaberin_3_-Fe^3+^_2_ bound to FoxA. Side view (top) and bird’s-eye view (bottom) are shown. **i,** Binding pocket interactions of bisucaberin_3_-Fe^3+^_2_ (orange) with FoxA (left) and overlay with FoxA-bound nocardamine-Fe^3+^ (orange; right) are shown. Nocardamine-Fe^3+^ structure from PDB:6Z8A was used for this comparison.

We reasoned that if bisucaberin is taken up by FoxA, it might compete with thiocillin for the TBDT, and tested this hypothesis using checkerboard assays with Δ*pvdA* Δ*pchA* Δ*foxA* expressing FoxA *in trans* (Fig. 5c). As expected, bisucaberin-Fe^3+^ antagonised thiocillin susceptibility. FoxA point mutations that reduced thiocillin uptake also negatively impacted growth of the Δ*pvdA* Δ*pchA* Δ*foxA* mutant supplemented with bisucaberin (Fig. 5d). Compared to the strain expressing WT FoxA, those expressing P659L or W503S required at least a 4-fold increase in bisucaberin to restore growth. These results further highlight the differences between the ligands, although there may be some overlap between the molecular determinants for thiocillin and bisuaberin uptake.

Since the FoxA point mutations reduced growth in the presence of bisucaberin, we tested whether the mutations inhibited bisucaberin binding (Fig. 5e). Saturation was not obtained even with 100 μg/mL of bisucaberin_3_-Fe^3+^_2_, suggesting that it has a lower affinity for FoxA than nocardamine. Thiocillin pre-treatment reduced quenching, similar to nocardamine. When this assay was repeated for the FoxA point mutants, bisucaberin_3_-Fe^3+^_2_ could still quench fluorescence of all the mutant transporters, suggesting the lack of growth for some FoxA mutations may instead be due to reduced bisucaberin uptake (Extended Data Fig. 7,8).

Interestingly, bisucaberin stimulated expression from P*_foxA_* in Δ*pvdA* Δ*pchA*, suggesting that it triggers the same signaling cascade as nocardamine (Fig. 5f). To confirm the promoter-GFP expression data, V5-tagged chromosomally-expressed mutants of WT FoxA and W503S were generated in Δ*pvdA* Δ*pchA* because they could support growth of the Δ*pvdA* Δ*pchA* Δ*foxA* mutant *in trans* (Fig. 5d, 5g). The mutants were treated with 5.0 μg/mL bisucaberin and examined for changes in FoxA expression. P659L T755V5 was excluded since it was not detectable (Fig. 2f). Both WT and W503S responded to bisucaberin treatment with increased FoxA expression (Fig. 5g).

To determine if molecular interactions of bisucaberin with FoxA resemble those of nocardamine, we determined, at 2.5Å resolution, the crystal structure of bisucaberin bound to FoxA (PDB: 8B43) (Fig. 5h, Extended Data Fig. 9). Bisucaberin_3_-Fe^3+^_2_ binds FoxA in a C-shape, unlike its linear configuration in the unbound form (Fig. 5i)^31^. Like nocardamine and ferrioxamine B, bisucaberin_3_-Fe^3+^_2_ forms a H-bond with Y805. Rings 2 and 3 of the bisucaberin complex contact the plug and barrel domains whereas ring 1 protrudes outwards and forms a second H-bond with Y664 (L8). A third H-bond is formed between ring 3 and Y799. The structural data agreed with the site-directed mutagenesis data – FoxA Y664A allows growth with nocardamine but not bisucaberin. H374, which forms a H-bond with nocardamine-Fe^3+^ is in the same binding pocket as the bisucaberin complex. H374Y may displace bisucaberin_3_-Fe^3+^_2_ in the binding pocket, reducing its interactions with other side chains. Nocardamine-Fe^3+^ from PDB:6Z8A was then overlaid with bisucaberin_3_-Fe^3+^_2_ to determine if there were differences in their interactions with FoxA (Fig. 5i). Interestingly, rings 2 and 3 of bisucaberin closely mimic the position of nocardamine, with the iron atoms superimposing one another. The hydroxamate moieties involved in chelation are also in similar orientations. Overall, the bisucaberin_3_-Fe^3+^_2_ complex binds FoxA in a similar position as nocardamine-Fe^3+^; but forms different H-bonds, and ring 1 interacts with L8, which is not seen with nocardamine. These data help to explain how thiocillin might interact with FoxA despite being larger than nocardamine.

### Sequence conservation of L8 is low and the loop is important for ligand uptake

Previously we tested a panel of Gram-negative bacteria capable of taking up nocardamine for their potential susceptibility to thiocillin. However, thiocillin showed only narrow spectrum activity against *P. aeruginosa* and some related Pseudomonads^28^. Since the data presented here show that single point mutations in FoxA were sufficient to confer thiocillin resistance, a phylogeny approach was used to determine whether the identity of key residues identified by the mutagenesis screen could explain resistance in other Gram-negatives. We constructed a phylogenetic tree for FoxA using shoot.bio^40,41^ in 606 bacterial species (Fig. 6a) and identified 16 orthologs^40^. Their sequences were aligned and residues of interest were examined for similarity to FoxA_PA14_ (Fig. 6b). Y374 was present in 6/16 orthologs while another 6 had H374, which we showed reduces nocardamine-Fe^3+^ binding and prevents thiocillin uptake (Fig. 2c,d). There was greater variation at positions corresponding to W503, K657, and Y664; however, some orthologs had S503, A657, and A664, which we showed compromise thiocillin binding and uptake (Fig. 2c,d). Additional substitutions at key residues were tested, and all mutations reduced thiocillin susceptibility without compromising nocardamine uptake (Fig. 6c,d). These results indicate that single amino acid differences are sufficient to confer resistance to thiocillin and may help to explain its narrow spectrum of activity against Gram negatives.

**Fig. 6.**
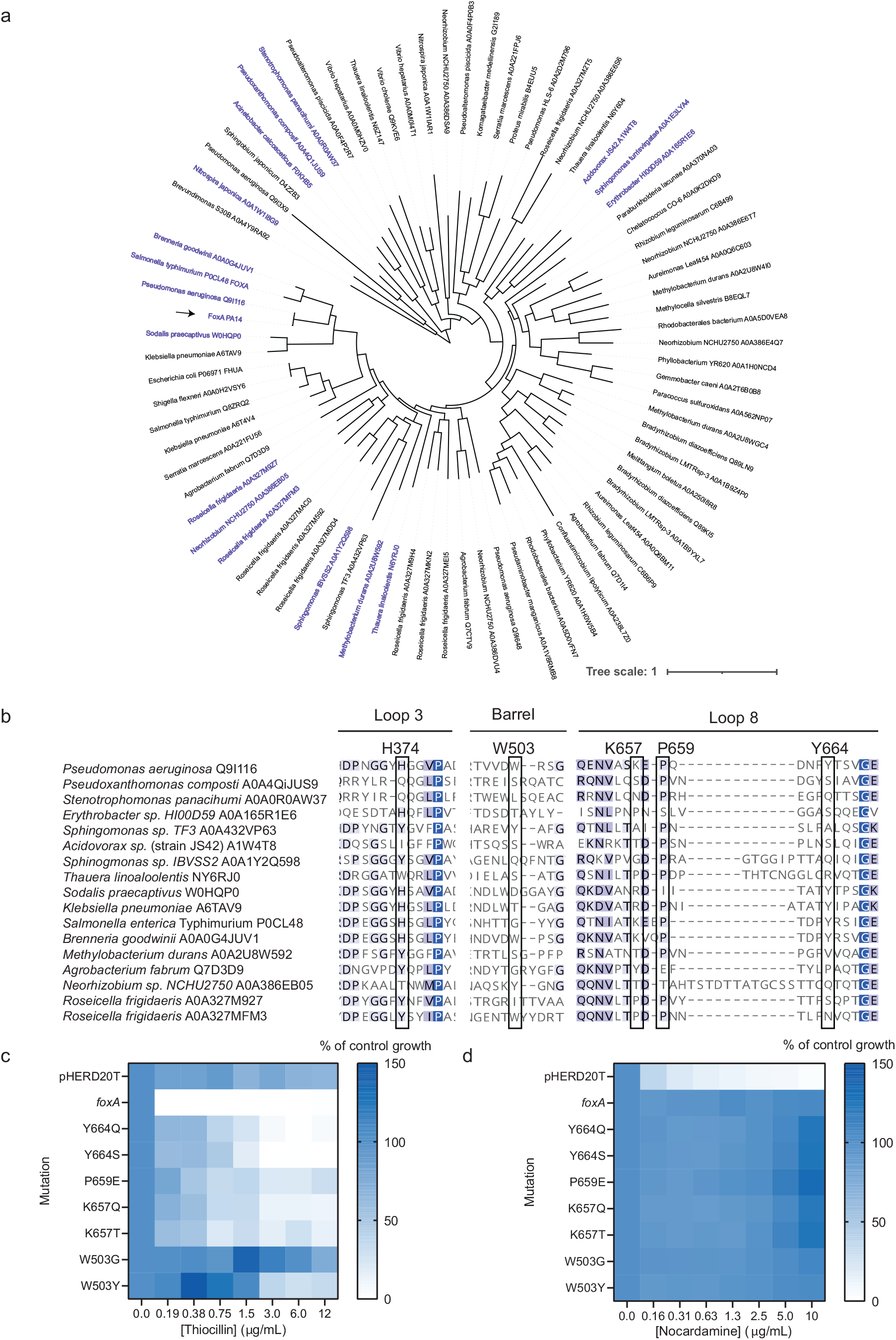
The amino acid sequences of FoxA orthologs vary at residues important for thiocillin susceptibility. **a,** Phylogenetic tree based on the PA14 FoxA amino acid sequence submitted to shoot.bio. FoxA_PA14_ is indicated by the arrow and orthologs are highlighted in purple. **b,** Alignment of FoxA orthologs with FoxA_PA14_. Residues important for thiocillin activity are boxed with the residue number corresponding to FoxA_PA14_ above. The intensity of purple indicates the degree of conservation of a particular amino acid residue with all orthologs where a darker shade indicates that most or all orthologs have a similar residue whereas white indicates that all or most orthologs have a different amino acid at the particular residue. **C, d**, PA14 Δ*foxA* expressing mutant FoxA from pHERD20T in 10:90 medium + 1% arabinose with increasing concentrations of thiocillin or (D) CAA + 1% arabinose with increasing concentrations of nocardamine.

## Discussion

In this study, we used molecular docking and site-directed mutagenesis to identify amino acid residues in FoxA necessary for thiocillin susceptibility. The results revealed that thiocillin and nocardamine interact differently with the TBDT despite occupying the same binding pocket. We focused on the potential role of L8, an extracellular loop that folds inwards when nocardamine binds^13^ because mutations in this loop reduced thiocillin uptake and activity. Similar loop rearrangements are observed with other transporters including FecA, indicating that this is a common feature of TBDTs^24^. Paradoxically, L8 was dispensible for thiocillin susceptibility – as seen with FoxA_ΔK657-T665_ – but essential for nocardamine uptake. These data expand on previous studies on the effects of loop deletions in FepA, which recognizes the siderophore enterobactin and the bacteriocin colicin B^42^. Deletion of FepA L2 was detrimental for both enterobactin uptake and colicin activity, indicating a similar transport mechanism, at least at the extracellular interface^42^. Our observations with thiocillin and nocardamine suggests that molecules do not need to mimic the native ligand in terms of structure or binding to use the same TBDT^30^. This observation is further supported by the lack of fluorescence quenching by thiocillin compared to nocardamine. Other studies showed that the bacteriocin, pyocin S2, mimics binding of pyoverdine^22^ with similar findings for the rifamycin derivative CGP 4832 and FhuA^43^, and the lasso peptide microcin J25 and FhuA^24^. Based on the thiocillin-FoxA docking predictions, L8 may be unable to fold because thiocillin sterically hinders this conformational change.

GFP-promoter and protein expression assays showed that L8 also contributes to signal transduction in a ligand-dependent manner, as nocardamine and bisucaberin stimulated FoxA expression while thiocillin did not. Although nocardamine uptake was supported by all single residue mutant transporters tested, the K657A and H374Y mutations compromised signaling. This result is consistent with prior studies showing that uptake of pyoverdine by FpvA and signalling could be separated^44^, suggesting they are independent processes. However, we found overlap between FoxA residues important for thiocillin and bisucaberin uptake and signal transduction by nocardamine, suggesting a series of conformational rearrangements not yet fully understood. Such connectivity may explain how thiocillin can be taken up through FoxA despite its structural differences from the native ligand. Similarly, bisucaberin uptake relied on residues similar to those required by thiocillin (Fig. 5d).

Overall, this work demonstrated that there are at least three ligands recognized by FoxA that interact with the TBDT in unique ways that yield different outcomes (Extended Data Fig. 7). These results highlight our as-yet incomplete understanding of the repertoire of ligands potentially recognized by different TBDTs; we suspect that many more molecules can use these transporters to cross the outer membrane. The structural complexity of natural product antimicrobials may relate to their evolution for uptake, in addition to target recognition. Discovery of such molecules, and understanding how they interact with the transporters and the consequences of those interactions, are important for guiding drug discovery and design. If an antimicrobial induces the expression of the transporter it uses to enter cells, it would potentiate its own uptake, an important consideration for drug development.

## Online Methods

### Strains, Media, and Compounds

All strains are listed in Supplementary Data Table 1. Lysogeny broth (LB) premixed powder, gentamicin, arabinose, and glucose were purchased from Bioshop. 10:90^27,28^ and CAA^6^ were made as previously described. Arabinose and glucose were prepared as a 20% solution in DI H_2_O or 10:90 and sterilized by filtration through 0.2μm membranes (Fisherbrand). Thiocillin, apo-nocardamine, nocardamine-Fe^3+^, bisucaberin, enterobactin, ferrichrome, and fluorescein-5-maleimide were purchased from Cayman Chemicals. Deferasirox was purchased from AK Scientific. Arthrobactin was purchased from MolPort. AlexaFluor 594 C5-maleimide was purchased from ThermoFisher Scientific. All compounds were dissolved in dimethyl sulfoxide and stored at −20°C.

### Docking of ligands into FoxA

SwissDock (http://www.swissdock.ch/)^33^, AutoDock VINA^32^, and HADDOCK (https://wenmr.science.uu.nl/haddock2.4/)^34^ were used to dock thiocillin into FoxA. A 3D structure of thiocillin was generated using Chem3D after energy minimalization. Docking by SwissDock was conducted using the web server where the target (FoxA) and the ligand (thiocillin) were submitted as PDB and mol2 files respectively. Two hundred and thirteen poses were generated and the pose with the lowest ΔG value (−10.9 kcal/mol) was selected to model binding. For AutoDock VINA, thiocillin was drawn in Chemdraw and transferred to Chem3D for energy minimalization prior to docking. Docking was conducted with the following grid parameters that encompasses the expected binding site and the surface of the plug: center_x = −9.628, center_y = 33.621, center_z = 5.268, size_x = 48, size_y = 42, size_z = 120. The pose with the lowest RMSD (0.000) and highest affinity (−10.5 kcal/mol) was selected to model binding. For HADDOCK, thiocillin (ligand) and FoxA (protein) were submitted as PDB files with the protein-ligand option selected and thiocillin indicated as a cyclic peptide. 191 structures were generated and grouped into three clusters. The largest cluster contained 177 structures and the structure with the best score was used for modeling.

### Minimal inhibitory concentration (MIC) assays

MIC assays were conducted as previously described^27,28^. Overnight cultures were grown in LB from a glycerol stock at 37°C with shaking (200rpm). Subcultures in either 10:90 or CAA (1:100 dilution) were cultured for four hours then adjusted to an OD_600_ of 0.1/500 in either fresh 10:90 or CAA. All compounds were serially diluted 2-fold in DMSO at 75x the final concentration. Plates were sealed to prevent evaporation and incubated at 37°C overnight in a shaking incubator (200rpm). The next day, the OD_600_ was determined with a plate reader (Thermo Scientific) and normalized to percent of growth of the vehicle control (DMSO) after subtracting the OD_600_ from blank media.

### Checkerboard assays

Checkerboards (8 rows x 8 columns) were conducted as previously described in a 96-well plate (Nunc)^27,28^. Thiocillin at 75x the final concentration dissolved in DMSO was added from bottom to top in increasing concentration. Siderophores at 75x the final concentration dissolved in DSMO was added from left to right in increasing concentration. Four columns were left for vehicle controls (DMSO) and sterile controls. Media with bacteria as described in the MIC assays were added to obtain a final volume of 150μL. Plates were sealed to prevent evaporation and incubated at 37°C overnight in a shaking incubator (200rpm). The next day, the OD_600_ was determined with a plate reader (Thermo Scientific) and normalized to percent of growth of the vehicle control (DMSO) after subtracting the OD_600_ from blank media.

### Molecular Biology

All primers are listed in Supplementary Data Table 2. All chromosomal mutants were generated by allelic exchange^45^. Primers flanking the upstream (^~^1000 bp) and downstream (^~^750 bp) regions of each gene of interest were amplified from PA14 genomic DNA (Promega Wizard Genomic DNA Purification Kit) and extracted with GeneJet Gel Extraction Kit (ThermoFisher). The upstream and downstream regions were joined by overlap extension PCR^46^ or three-piece ligations, digested with the indicated enzymes (FastDigest ThermoFisher), and ligated into pEX18Gm to make each deletion construct (T4 DNA ligase, ThermoFisher). The ligation mixtures were transformed into competent *E. coli* DH5α by heat shock with a recovery period of three hours in LB. Cells were plated on LB 1.5% agar containing 15μg/mL gentamicin supplemented with X-gal for blue-white screening. The plates were incubated at 37°C overnight. White colonies were selected and grown in LB + 15μg/mL gentamicin. Plasmids were isolated using the GeneJet Plasmid Miniprep Kit (ThermoFisher).

Plasmids with correct inserts were transformed into competent *E. coli* SM10 by heat shock with a recovery period of 3 h in LB. The cells were plated on LB 1.5% agar containing 15μg/mL gentamicin and grown overnight at 37°C. One colony was inoculated in LB + 15μg/mL gentamicin overnight. The strain of interest was also inoculated from a single colony into LB. Both cultures were grown overnight at 37°C. SM10 with the desired deletion construct was mated with PA14 by mixing 500 μL of each overnight culture in a 1.5mL centrifuge tube. Cells were spun down and the supernatant was removed. Cells were resuspended in 50 μL fresh LB, spotted on LB 1.5% agar, and incubated overnight at 37°C. Cells were streaked onto *Pseudomonas* Isolation Agar (PIA, Difco) supplemented with 100μg/mL gentamicin and incubated overnight at 37°C. Single colonies were streaked onto LB + 15% sucrose (BioShop) and incubated overnight at 37°C. To check for colonies with the correct deletion, 16 colonies were patched onto LB + 15% sucrose and LB + 30μg/mL gentamicin and incubated overnight at 37°C. Patches that grew on the sucrose plates but not gentamicin plates were checked by colony PCR with primers flanking the deleted gene and internal primers and compared to WT controls. Patches with the desired deletions were streaked onto LB + 15% sucrose to isolate single colonies, incubated overnight at 37°C and checked again by colony PCR. A single colony was inoculated into LB broth and the process was repeated to generate double and triple mutants.

Overexpression strains were generated with the plasmid pHERD20T, an arabinose-inducible expression vector with the P_BAD_ promoter under control of AraC. Primers flanking the gene of interest including the ribosome binding site were amplified from *P. aeruginosa* PA14 genomic DNA, digested with the desired restriction enzymes, and ligated into pHERD20T digested with the same enzymes. Ligation mixtures were added to chemically competent DH5α and heat shocked with a recovery period of 1-2 hours in LB at 37°C. All the cells were plated on LB 1.5% agar supplemented 100μg/mL ampicillin, X-gal, and arabinose and incubated at 37°C overnight. White colonies were checked by colony PCR and colonies with correct plasmids were cultured in LB broth supplemented with 100μg/mL ampicillin. Plasmids were isolated, checked for the correct insert by restriction digest, and electroporated into the desired strain or mutant with a recovery of 1-2 hours in LB at 37°C. All the cells were plated on LB 1.5% agar supplemented with 200μg/mL carbenicillin (AK Scientific). A single colony was picked and grown in LB broth supplemented with 200μg/mL carbenicillin overnight at 37°C and used to make glycerol stocks and for subsequent assays. Correct inserts were verified by Sanger sequencing by the McMaster Genomics Facility Mobix Lab.

### Cysteine labeling, fluorescence quenching, and recovery assays

PA14 Δ*foxA* pHERD20T-*foxA* cysteine mutants were subcultured 1:100 from LB overnight cultures into 10:90 + 2% arabinose. Cultures were grown for four hours at 37°C with shaking (200 rpm). Cells were harvested by centrifugation (6000 x g, 5 min) and resuspended in 1mL of PBS (or PBS + 0.4% glucose for uptake studies) + 10μM fluorescein-5-maleimide (Cayman Chemicals) or AlexaFluor 594 C5 maleimide (ThermoFisher) for microscopy. A 1X PBS working buffer was prepared from a 10X stock (80g NaCl, 2g KCl, 26.8g Na_2_HPO_4_-7H_2_O, 2.4g KH_2_PO_4_ in 1L of de-ionized H_2_O, pH 7.4). Cells were labeled at 37°C with shaking (200rpm) for 30 min in the dark. Excess dye was quenched with 1 mM dithiothreitol (DTT; Sigma) to stop the reaction. Cells were washed 3X with PBS. For fluorescence recovery assays, cells were incubated with 1X PBS + 0.4% glucose at 37°C with shaking at 200 rpm. Glucose was prepared as a 20% stock (wt/v) in 1X PBS and filter sterilized. Cells were spun down and washed 3X with 1X PBS and resuspended in 1X PBS for quenching assays and 1X PBS + 0.4% glucose for fluorescence recovery assays. Cells were distributed into black 96-well plates (Corning) with 146 μL of OD_600_ 0.1 cells per well. For pre-treatment with and without thiocillin, 2μL of thiocillin (final concentration of 12μg/mL – equivalent to 10μM) or DMSO was added to the cells and incubated at room temperature for 5 min before 2 μL of nocardamine-Fe^3+^ or bisucaberin-Fe^3+^ was added. Fluorescence of fluorescein-5-maleimide (ex. 494nm em. 520nm) was read during the thiocillin pre-treatment period and immediately after nocardamine-Fe^3+^ or bisucaberin-Fe^3+^ was added (BioTek Neo). Fluorescence recovery assays were conducted as above but plates were incubated at 37°C and fluorescence was recorded every minute for one hour. Since thiocillin reduced quenching compared to nocardamine-Fe^3+^ alone, differences in baseline quenching were normalized by subtracting F/F_0_ at t = 5 min, the time at which nocardamine-Fe^3+^ was added, from F/F_0_ at each timepoint to obtain ΔF/F_0_.

### Microscopy

AlexaFluor594 C5-maleimide-labeled cells (5μL) were spotted onto a 1% agarose pad on a microscope slide. The agarose pad was mounted with a glass coverslip directly prior to imaging. Cells were imaged using brightfield and fluorescence microscopy on a Nikon A1 confocal microscope through a Plan Apo 60X (NA=1.40) oil objective. Image acquisition was done using Nikon NIS Elements Advanced Research (Version 5.11.01 64-bit) software. Cells were imaged using an EVOF FL Auto microscope through a Plan Apo 60X (NA=1.40) oil objective.

### Promoter assays

P*_cdrA_-gfp*^47^ was purchased from Addgene. P*_cdrA_* was removed by digestion with XbaI and SacI followed by ligation of P*_foxA_* with the same synthetic ribosome binding site into the vector (Supplementary data). The plasmid was introduced into PA14, PA14 Δ*pvdA* Δ*pchA*, and PA14 FoxA mutants by electroporation. Siderophores (apo-nocardamine, apo-bisucaberin) and thiocillin were added to cells the same way MIC assays are conducted in Corning black 96-well plates with clear bottoms. After incubation overnight at 37°C with shaking (200rpm), GFP production (excitation: 494nm and emission: 515nm)^47^ and OD_600_ were determined (BioTek Neo).

### Outer membrane preparations

PA14 mutants and strains were grown overnight in LB with 200μg/mL carbenicillin (for pHERD20T strains) or without (for chromosomal mutants) at 37°C with shaking (200 rpm). Cells were subcultured 1:100 into 10:90 or CAA for 4 h then diluted to OD_600_ 0.1/500 in 50mL of fresh 10:90 (with or without arabinose for plasmid-carrying strains and chromosomal mutants) or CAA (for chromosomal mutants). Cultures were grown overnight at 37°C with shaking (200 rpm). Cells were harvested by centrifugation and resuspended in 10 mM Tris pH 8.0.

Cells were lysed by sonication (Misonix Sonicator 3000) on ice (30 s pulse, 30 s rest, 3 min, power level 5.0). Cell debris was removed by centrifugation (6000 x g, 5 min, 4°C). Membranes were harvested at 21000 x g for 30 min at 4°C. The pellet was resuspended in 100 μL de-ionized H_2_O and combined with 900μL 11.1mM Tris 3% sarkosyl pH 7.6 and incubated at room temperature for 30 min with shaking (200rpm). Insoluble proteins were collected by centrifugation at 21000 x g for 30 min at 4°C.

### SDS-PAGE and Western Blots

Outer membrane preparations were resuspended in 20 μL 1X loading buffer for SDS-PAGE analysis. Each lane was loaded with 10 μL of outer membrane preparation and separated at 80 V for 15 min and 120 V for 1 h. Gels were transferred to nitrocellulose membranes (225 mA, 1 h) and blocked with 5% skim milk in PBS pH 7.4 overnight at 4°C. Primary antibodies (rabbit α-V5 (Sigma 21160752), mouse α-DYKDDDDK (Invitrogen MA1-91878), and rabbit #3198 α-PilF) were used at 1:1000 dilutions in PBS and incubated with the blot at room temperature for 1 h. Blots were washed 3x with PBS and incubated with goat α-rabbit-alkaline phosphatase (AP) or rabbit α-mouse-AP for 1 h in PBS at 1:2000 dilutions. Blots were washed 3X with PBS before development with AP buffer (1mM Tris, 100mM NaCl, 5mM MgCl_2_ pH 9.5) + 5-bromo-4-chloro-3-indoyl phosphate (BCIP) + nitro-blue tetrazolium (NBT) for 15-30 min. Blots were imaged on an Azure C400 Imaging System.

### Thiocillin uptake assays, HPLC, and mass spectrometry

PA14 strains were grown overnight in LB with 200μg/mL carbenicillin at 37°C with shaking (200 rpm). Cells were subcultured 1:100 into 10:90 + 2% arabinose for 4 h and collected by centrifugation. Cells were washed 3X in 10:90 and resuspended in fresh 10:90 + 12μg/mL (10μM) thiocillin to an OD_600_ of 1.0. Cells were aliquoted into 1.5mL centrifuge tubes and incubated at 37°C with shaking (200rpm) for 1h then collected by centrifugation. Cells were washed 3x with PBS then resuspended in 100 μL H_2_O. Cultures were subjected to three cycles of freeze-thaw at −80°C and 60°C for 5 min per freeze-thaw. Lysed cells were collected by centrifugation for 10 min at 21000 x g at room temperature and the supernatant was collected in a clean centrifuge tube. Methanol (100 μL) was added to the cell pellet and mixed to collect residual thiocillin and centrifuged for 10 min at 21000 x g. The methanol layer was combined with the previous supernatant and mixed.

LC-MS was used to analyze the concentration of thiocillin in the samples (Eclipse Plus C18 column, 2.1 × 100mm, 1.8μm; solvent A: water + 0.1% formic acid; solvent B: acetonitrile + 0.1% formic acid; flow rate 0.3mL/min; gradient 95:5 A:B; total run time = 10 min; injection volume = 5 μL; MS method: QQQ; mode: positive). Matrix effects were accounted for by preparing thiocillin standards in blank samples from PA14 Δ*foxA* pHERD20T. Thiocillin concentrations were normalized based on the CFUs from each sample.

### Bisucaberin-FoxA co-crystallization and data collection

FoxA was overexpressed in *Escherichia coli* Lemo21 DE3 cells as previously described^13^. Cells were grown in 2xYT media supplemented with 0. 5 mM L-rhamnose at 37 °C to an OD_600_ of 0.8-1 and induced with 0.1 mM isopropyl β-D-1-thiogalactopyranoside. Cells were grown at 19°C for 16-18 h and lysed using high-pressure homogenizer (EmulsiFlex-C3, Avestin). Cell debris were removed by centrifugation at 22,000 g for 30 min, and supernatant was incubated with 1% Triton X-100 for 1 hr. The outer membrane fraction was isolated by centrifugation at 150,000 g for 1 h and solubilized using 1% octyl-β-glucoside and 1% C8E4 in TBS buffer (20 mM Tris-HCl pH 7.5, 200 mM NaCl). Solubilised outer membranes were cleared by another round of centrifugation at 150,000 g for 20 min and loaded onto Ni^2+^-NTA beads for 1 h. The beads were washed with TBS buffer supplemented with 30 mM imidazole and 0.6% C8E4 and eluted with 300 mM imidazole in the same buffer. Fractions containing the protein were pooled and tobacco etch virus protease (TEV, 10:1 w/w) was added overnight to cleave the His6-tag from FoxA. Next day, the eluate was passed through the Ni-NTA beads to remove TEV and any impurities, the flow-through concentrated to 500 μl and loaded onto a Superdex S200 10/300 size-exclusion column. Peak fractions were concentrated to 6-8 mg/ml and used for crystallization or stored at −80 °C.

Purified FoxA in C8E4 detergent was mixed with 10-fold excess Fe-bisucaberin, incubated on ice for 30 min and used for crystallization. Crystallisation conditions were 1.8 M ammonium sulfate, 0.1 M MES pH 6.5, 0.8% OG. Crystals typically appeared within 1-2 weeks at 19 °C.

All X-ray diffraction data were collected at PETRA III/EMBL P13 beamline, dataset processed with XDS^48^, merged with AIMLESS^49^, anisotropy correction was performed with STARANISO server. The structure was determined using molecular replacement with PHENIX Phaser using apo FoxA as a starting model (PDBID:6I98) and refined using phenix.refine^50^. Structural coordinates have been deposited in the RCSB Protein Data Bank under the accession number 8B43. Data collection and refinement statistics are found in Supplementary Data Table 3.

### Phylogenetic Tree, ortholog identification, and sequence alignments

The FoxA phylogenetic tree and ortholog identification was conducted using SHOOT^40^ (<u>shoot.bio</u>). The tree was visualized with iTOL^41^ (https://itol.embl.de/). Ortholog sequences were imported from Uniprot^51^ and sequences were aligned in Geneious Prime by MUSCLE alignment^52^. Accession numbers for the orthologs are as follows: *Pseudomonas aeruginosa* Q9I116, *Pseudoxanthomonas composti* A0A4QiJUS9, *Stenotrophomonas panacihumi* A0A0R0AW37, *Erythrobacter sp. HI00D59* A0A165R1E6, *Sphingomonas sp. TF3* A0A432VP63, *Acidovorax sp*. (strain JS42) A1W4T8, *Sphinogmonas sp. IBVSS2* A0A1Y2Q598, *Thauera linoaloolentis* NY6RJ0, *Sodalis praecaptivus* W0HQP0, *Klebsiella pneumoniae* A6TAV9, *Salmonella enterica* Typhimurium P0CL48, *Brenneria goodwinii* A0A0G4JUV1, *Methylobacterium durans* A0A2U8W592, *Agrobacterium fabrum* Q7D3D9, *Neorhizobium sp. NCHU2750* A0A386EB05, *Roseicella frigidaeris* A0A327M927, *Roseicella frigidaeris* A0A327MFM3.

## Acknowledgements

We thank Nikki Henriquez from the Centre for Microbial Chemical Biology (CMCB) at the Institute for Infectious Diseases Research (IIDR) for assistance with experiments to measure intracellular concentrations of thiocillin. We are grateful to the staff at beamlines P13 and P14 (EMBL, Hamburg) and acknowledge access to the Sample Preparation and Characterization (SPC) Facility of EMBL, Hamburg. This work was supported by a Natural Sciences and Engineering Research Council Discovery Grant RGPIN-2021-04237 to LLB, a Canadian Institutes of Health Research grant (FRN-148463) to GDW, and the excellence cluster ‘The Hamburg Centre for Ultrafast Imaging - Structure, Dynamics and Control of Matter at the Atomic Scale’ of the Deutsche Forschungsgemeinschaft (DFG EXC 1074) to IJ and HT. LLB and GDW hold Tier 1 Canada Research Chairs in Microbe-Surface Interactions and Antimicrobial Biochemistry, respectively. DCKC holds a Canadian Institute of Health Research (CIHR) Canada Graduate Scholarship – Doctoral program (CGS-D).

## Author Contribution

DCKC and LLB designed the experiments and wrote the draft manuscript. KK and GDW helped with method development for the thiocillin uptake study and KK performed mass spectrometry on initial samples. IJ isolated and purified FoxA for crystallization with bisucaberin. All authors provided editorial input on the final manuscript.

## Data availability

All data generated or analysed during this study are included in this published article and its Supplementary Information. Structural coordinates and structural factors have been deposited in the RCSB Protein Data Bank under accession number 8B43 (see Table S1, Suppl. Info.).

## EXTENDED DATA

**Extended Data Fig. 1.**
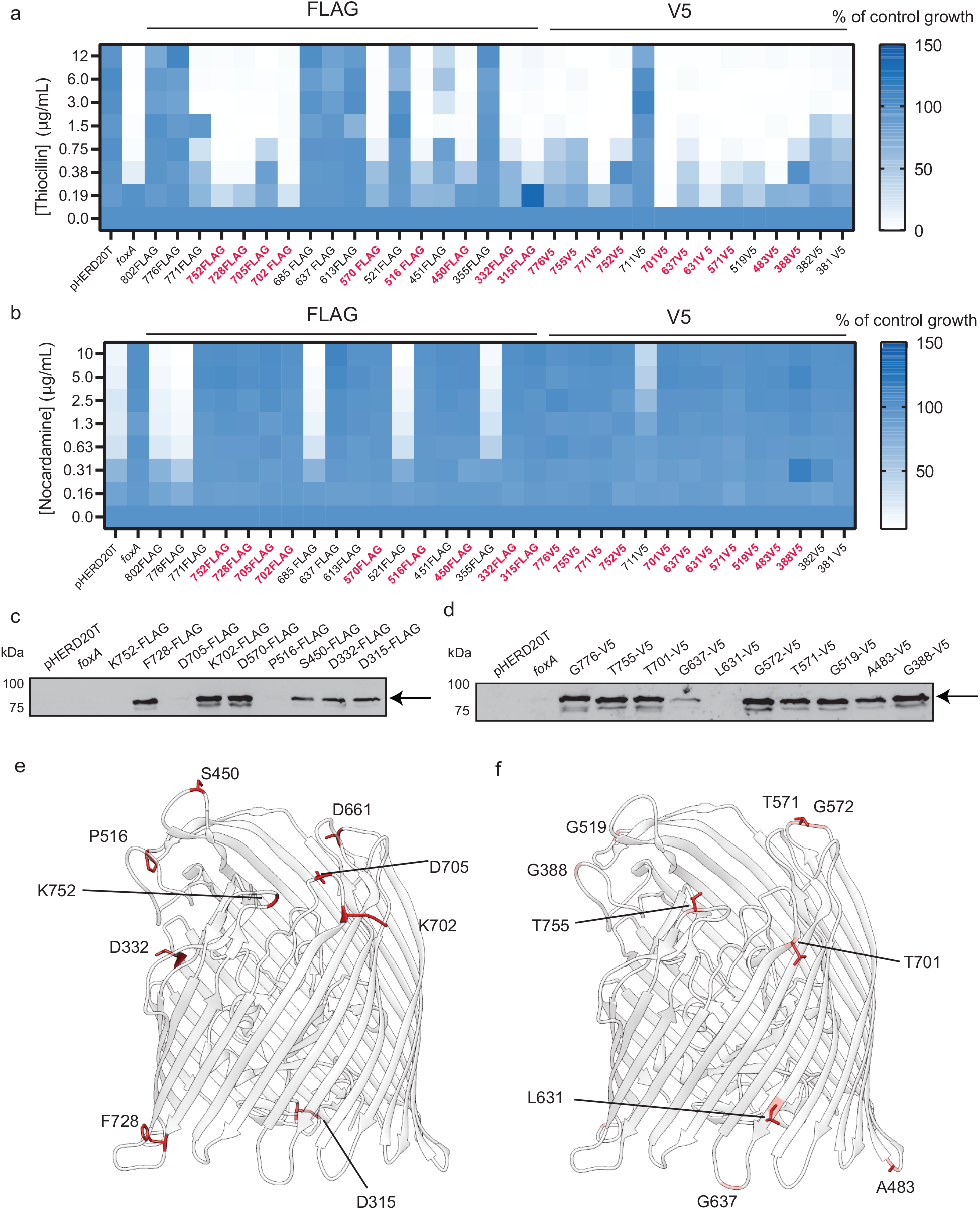
Screening for permissive tag locations on FoxA for insertion of FLAG or V5 epitopes. Epitope tags were introduced into *foxA* by overlap extension PCR and the resulting proteins expressed *in trans*. **A,** Thiocillin MIC assay with PA14 Δ*foxA* harbouring tagged *foxA* in pHERD20T tested in 10:90 + 1% arabinose. Transporters that resensitized Δ*foxA* to thiocillin are bolded in red. **B,** Nocardamine growth assay with PA14 Δ*foxA* harbouring tagged *foxA* in pHERD20T in 10:90 + 1% arabinose. Transporters that complemented Δ*foxA* growth with nocardamine are bolded and highlighted in red. **c,** Detection of selected FLAG tags in strains with WT susceptibility to thiocillin and that grow in the presence of nocardamine. Empty vector and WT FoxA were used as negative controls. **d,** Detection of selected V5 tags in strains that had WT susceptibility to thiocillin and that grow in the presence of nocardamine. Empty vector and WT FoxA were used as negative controls. **e** and **f,** Locations of inserted FLAG and V5 epitope tags respectively (PDB:6I98).

**Extended Data Fig. 2.**
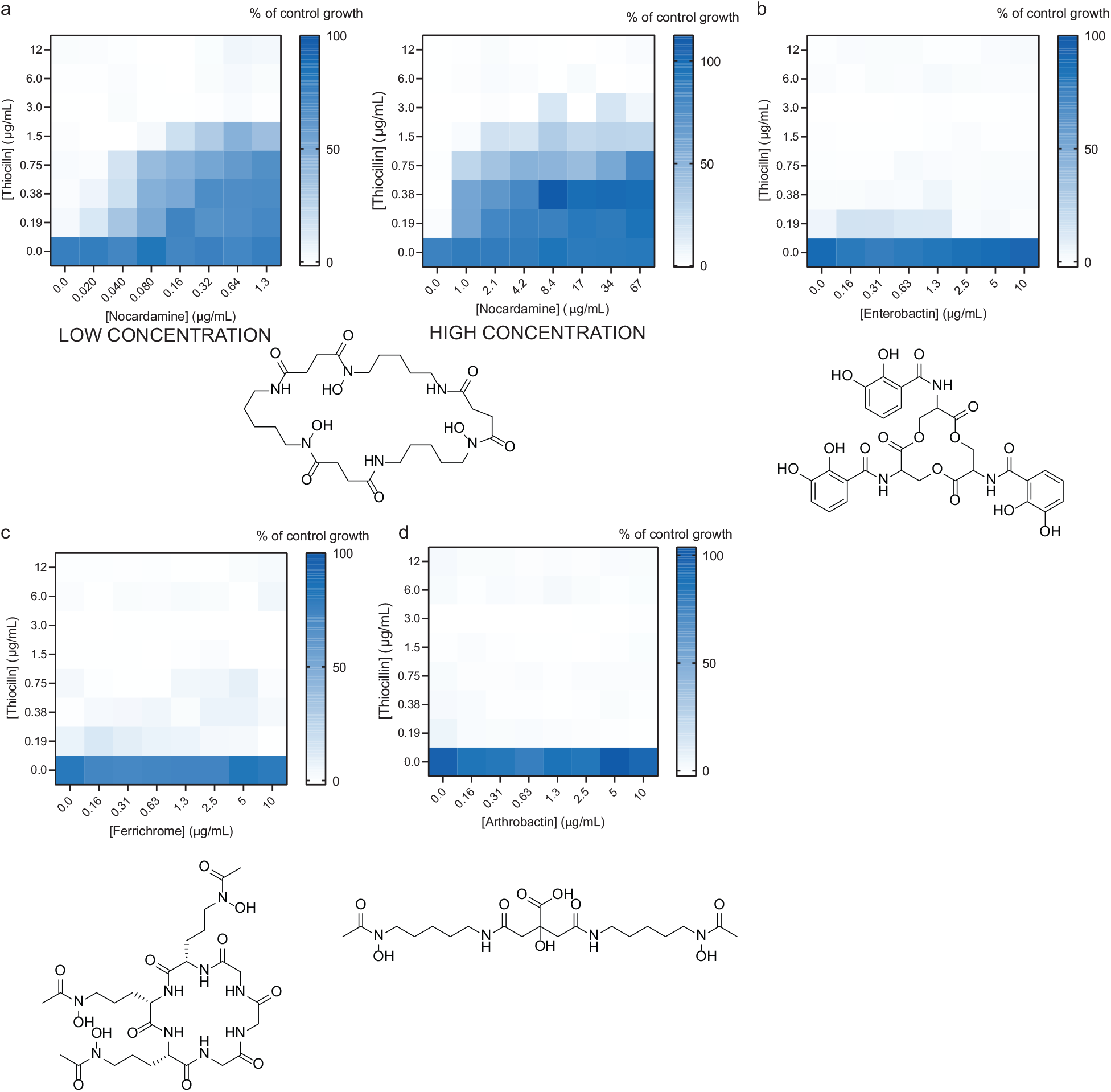
Only nocardamine antagonizes thiocillin uptake. Checkerboard assays of PA14 Δ*foxA* pHERD20T-*foxA* grown with thiocillin and different siderophores. **a,** Nocardamine (highest concentration tested at 1.3 μg/mL and 67 μg/mL), **b,** enterobactin, **c,** ferrichrome, and **d,** arthrobactin in 10:90 + 1% arabinose. Structures of the siderophores are shown. Growth is expressed as percent of vehicle control where blue indicates growth and white indicates no growth. All checkerboards are averages of three independent biological replicates.

**Extended Data Fig. 3.**
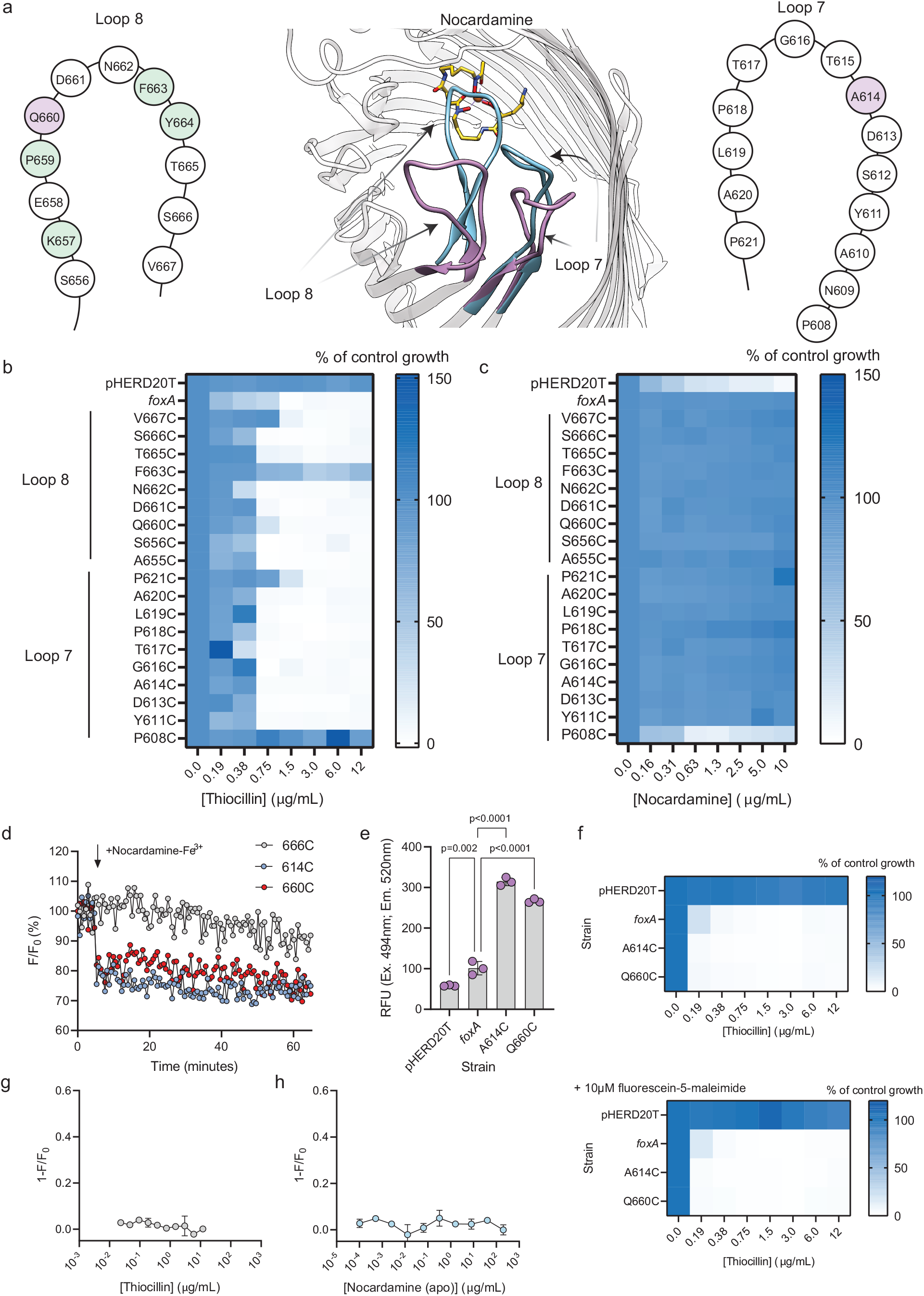
Screening for permissive cysteine-labeling sites for FoxA that allow WT responses to nocardamine. Cysteine mutagenesis of L7 and L8 was conducted to find residues that could be used for detecting nocardamine-Fe^3+^ binding. **a,** Depiction of L7 and L8 movement of FoxA upon siderophore binding generated from PBD 6I96 and 6I97, respectively. Loop orientation before and after siderophore binding are shown in purple and blue respectively. Ferrioxamine B-Fe^3+^ is shown in orange. L8 (left) and L7 (right) amino acid compositions are shown with residues important for thiocillin susceptibility highlighted in green. Residues that can be used for detection are highlighted in purple. PA14 Δ*foxA* expressing L7 and L8 cysteine mutants *in trans* were tested for susceptibility to **b,** thiocillin and **c,** ability to grow with nocardamine. Thiocillin tests were conducted in 10:90 + 1% arabinose while apo-nocardamine tests were conducted in iron-limited CAA + 1% arabinose. **d,** Screening of fluorescein-5-maleimide labeled FoxA cysteine mutants for fluorescence quenching upon addition of nocardamine-Fe^3+^. Fluorescence was measured for 5 min initially (Ex. 494nm Em. 518nm) and nocardamine-Fe^3+^ was added where indicated by the arrow. All cysteine mutants were screened with 0.050 μg/mL nocardamine-Fe^3+^. Fluorescence of labeled A614C and Q660C was quenched. S666C is shown as an example of the remaining transporters that did not respond to addition of nocardamine-Fe^3+^. **e,** Labeling by fluorescein-5-maleimide of A614C and Q660C compared to WT FoxA and empty vector controls. Three independent biological replicates are shown. One-way ANOVA followed by Dunnett’s test was used for statistical determinations. **, p<0.001; **** p<0.0001. **f,** Thiocillin activity in the absence (top) and presence of 10μM fluorescein-5-maleimide. Three independent biological replicates were conducted with the averaged result shown. **g,** Quenching by thiocillin and **h,** apo-nocardamine using the A614C sensor strain. Results shown are averaged from two independent biological replicates.

**Extended Data Fig. 4:**
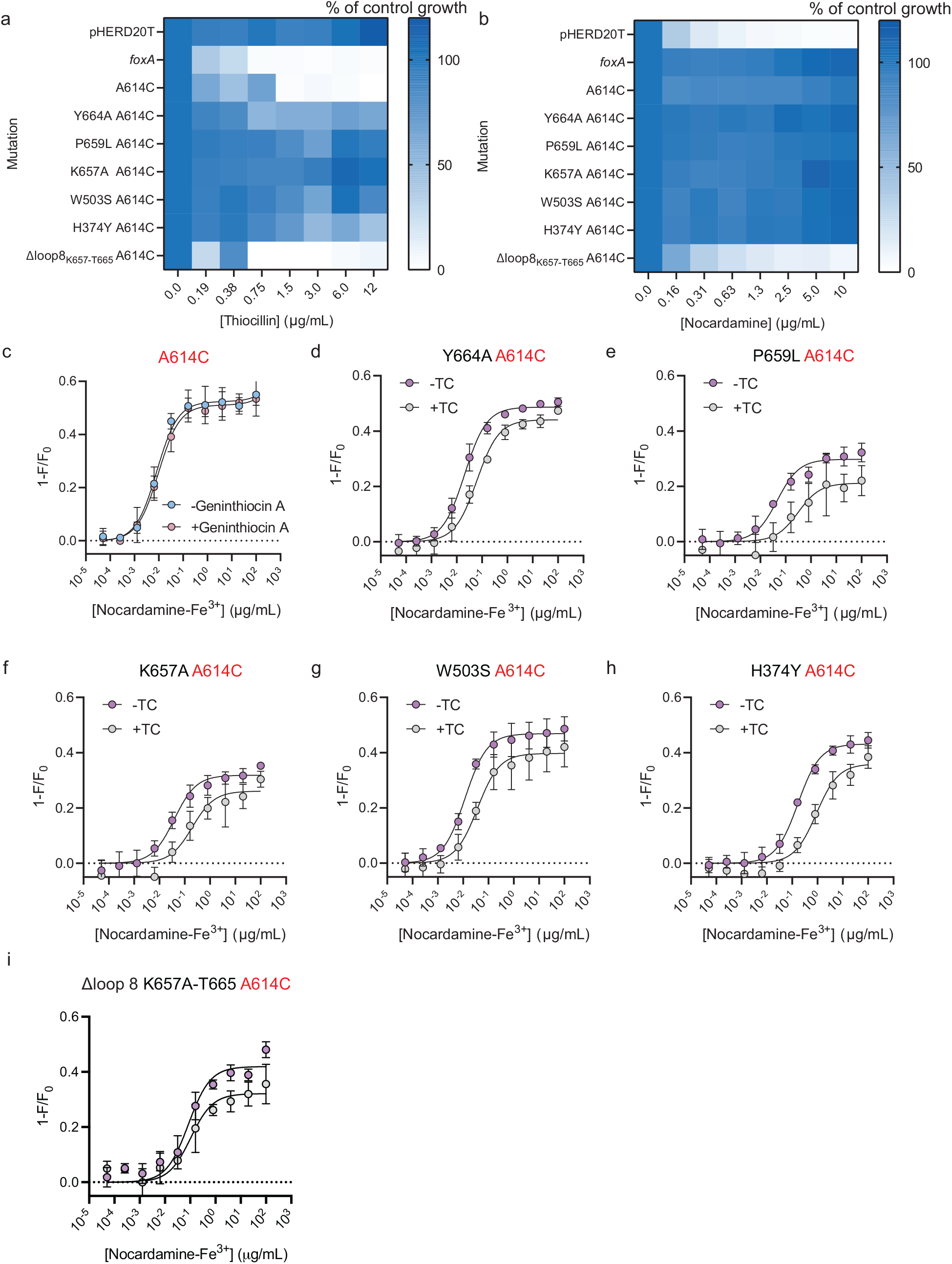
Labeled single residue mutants have reduced affinity for nocardamine-Fe^3+^ and addition of thiocillin further reduces the K_d_. **a,** MIC assays with PA14 Δ*foxA* expressing labeled single amino acid mutants treated with thiocillin. **b,** MIC assays with PA14 Δ*foxA* expressing labeled single amino acid mutants treated with apo-nocardamine. **c,** Fluorescence quenching of FoxA A614C labeled with fluorescein-5-maleimide. Cells were treated without (blue circles) or with 11 μg/mL (10 μM) geninthiocin A (red circles) for 5 mins prior to nocardamine-Fe^3+^ addition. **d-i,** Fluorescence quenching of fluorescein-5-maleimide-labeled FoxA mutants. Results are averaged from three independent biological replicates with standard deviations shown. Purple circles indicate quenching in the absence of thiocillin pre-treatment whereas grey circles indicate quenching after thiocillin pre-treatment.

**Extended Data Fig. 5.**
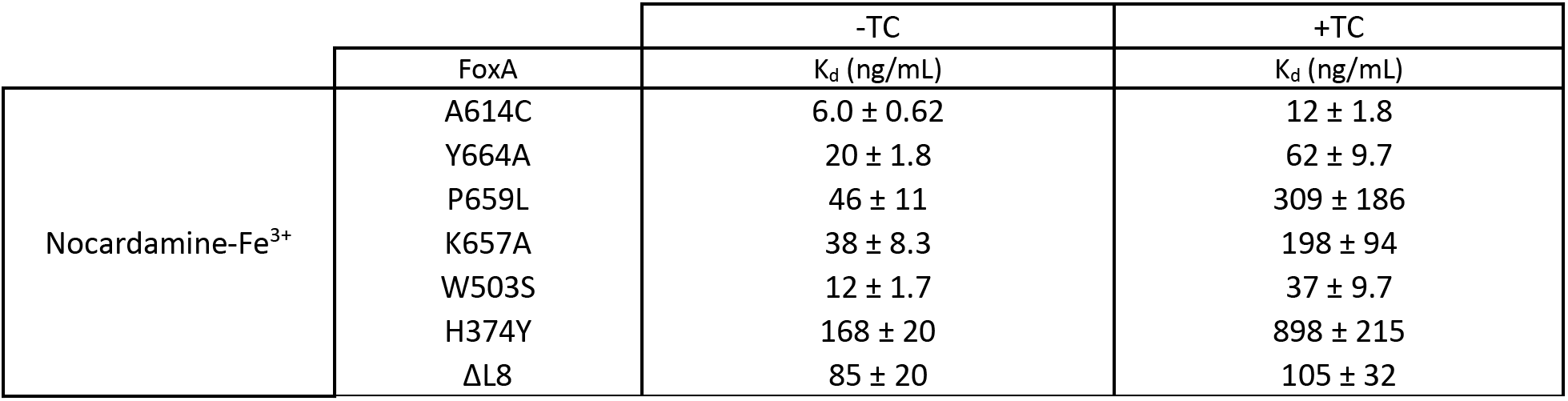
Summary of K_d_ values from WT and mutants treated with nocardamine and thiocillin. Values are reported as ng/mL averaged from three independent biological replicates.

**Extended Data Fig. 6.**
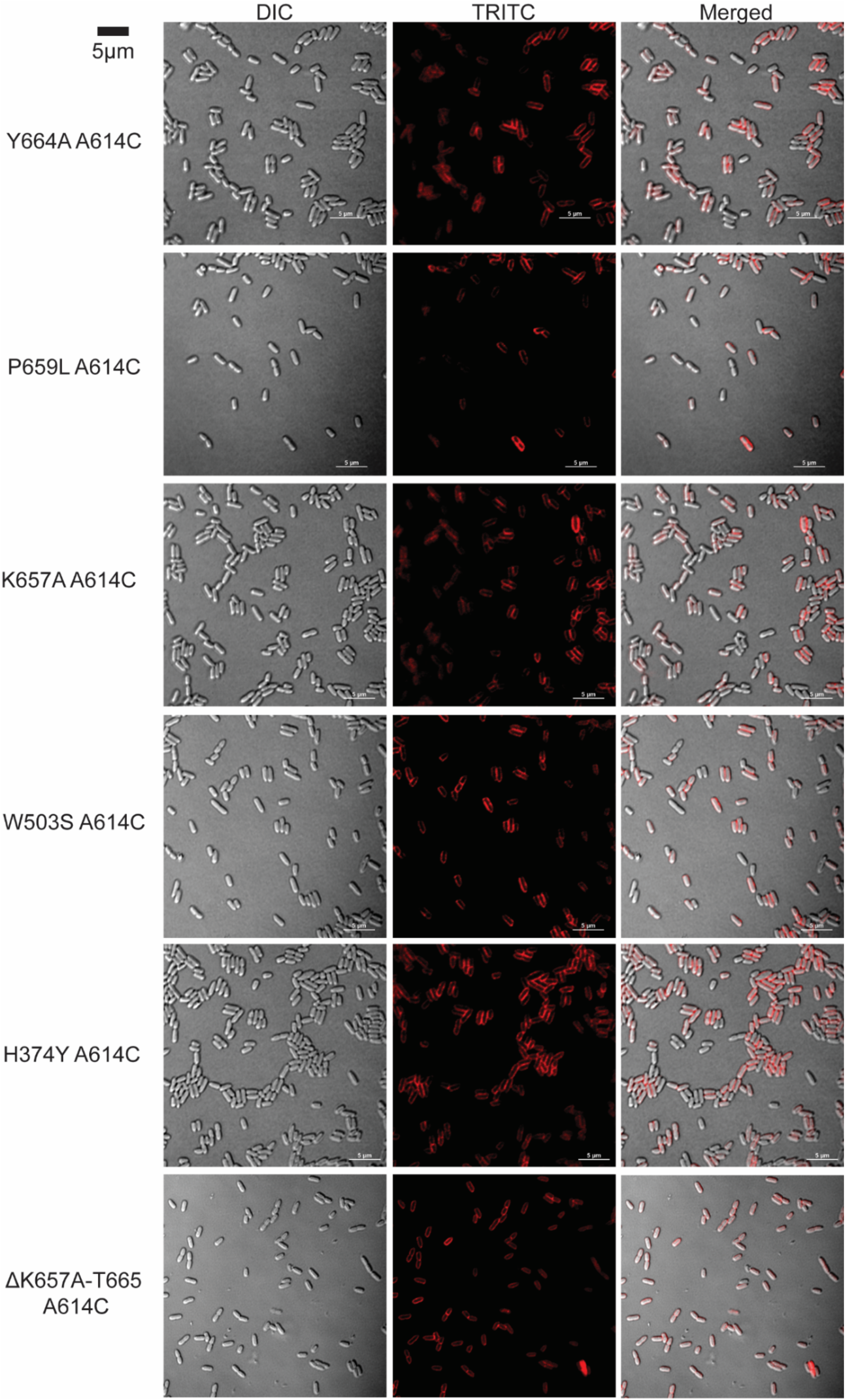
Confocal microscopy images of AlexaFluor 594 labeled FoxA A614C with amino acid mutations. Representative images for each strain are shown. Scale bar = 5μm.

**Extended Data Fig. 7.**
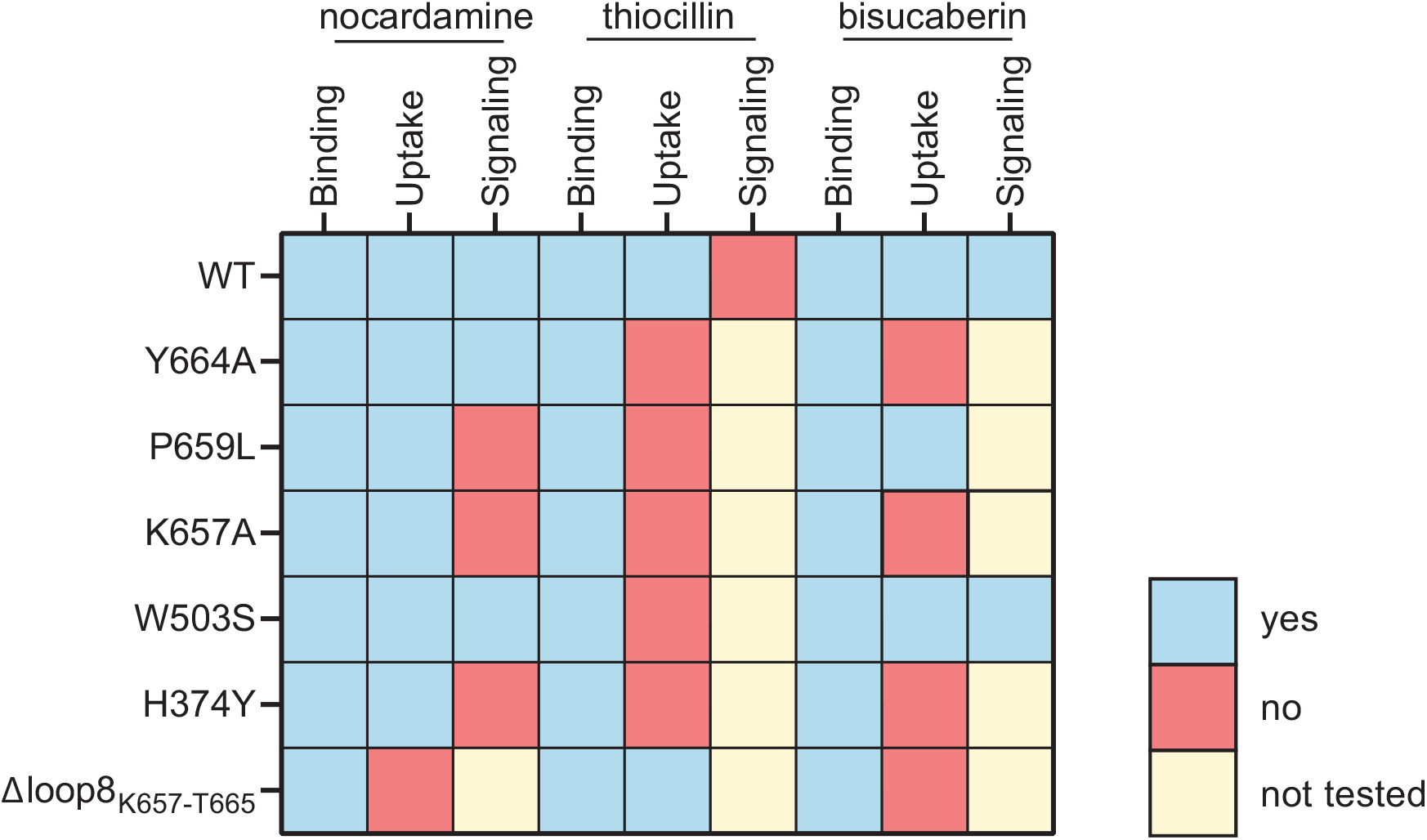
Summary of phenotypes of FoxA mutants. Mutations are listed on the left and their effects on ligand binding, uptake, and signaling are indicated. Blue boxes indicate that the phenotype was observed. Red boxes indicate that the phenotype was not observed or disrupted. Yellow boxes indicate that the phenotype was not tested for a particular mutation. If uptake was not observed, then signaling was not tested. Signaling for P659L was not tested since the V5 tag was not detected even *in trans*.

**Extended Data Fig. 8.**
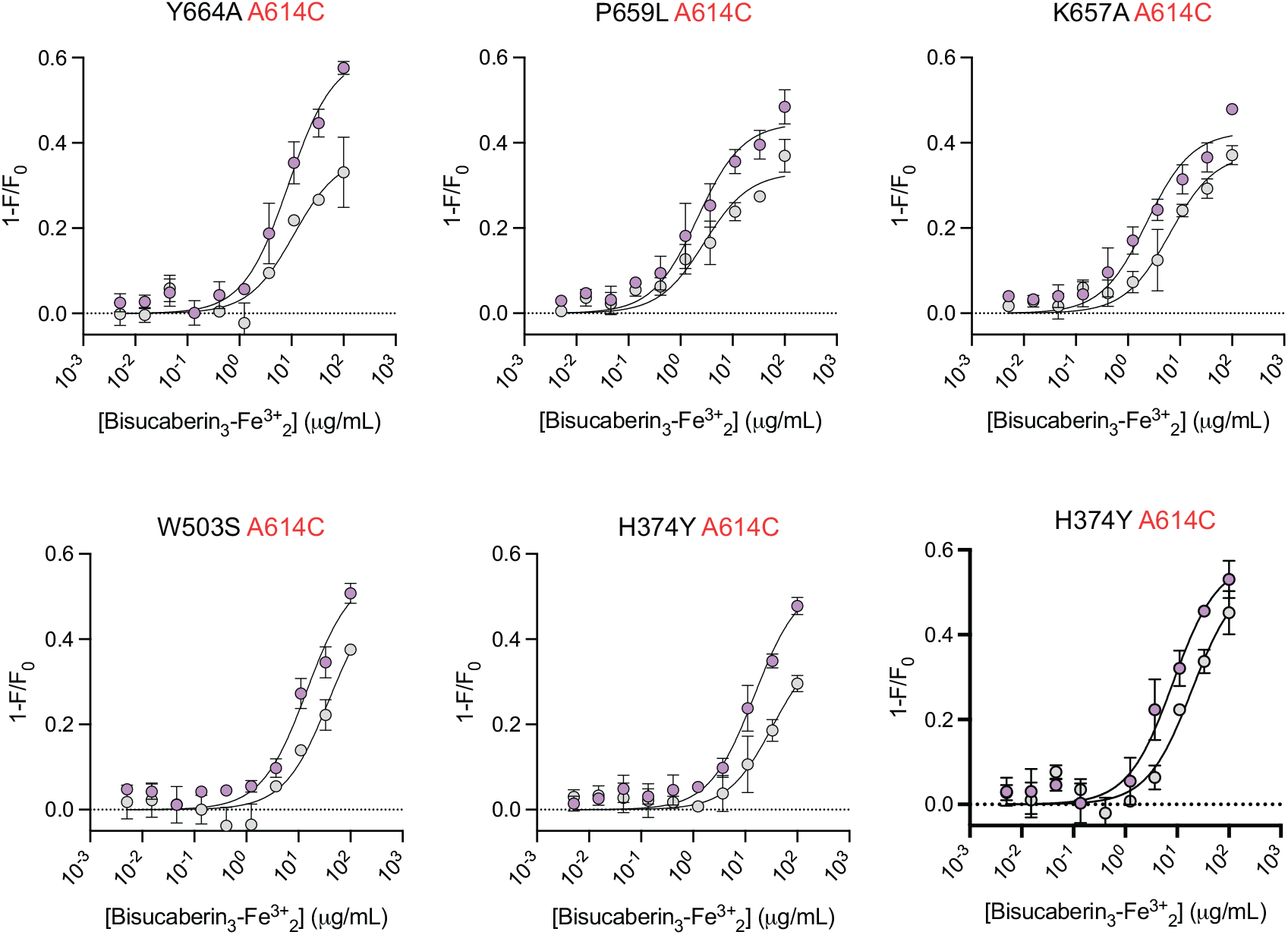
Fluorescence quenching of labeled FoxA and FoxA mutants by bisucaberin with and without thiocillin. Results are averaged from three independent biological replicates.

**Extended Data Fig. 9.**
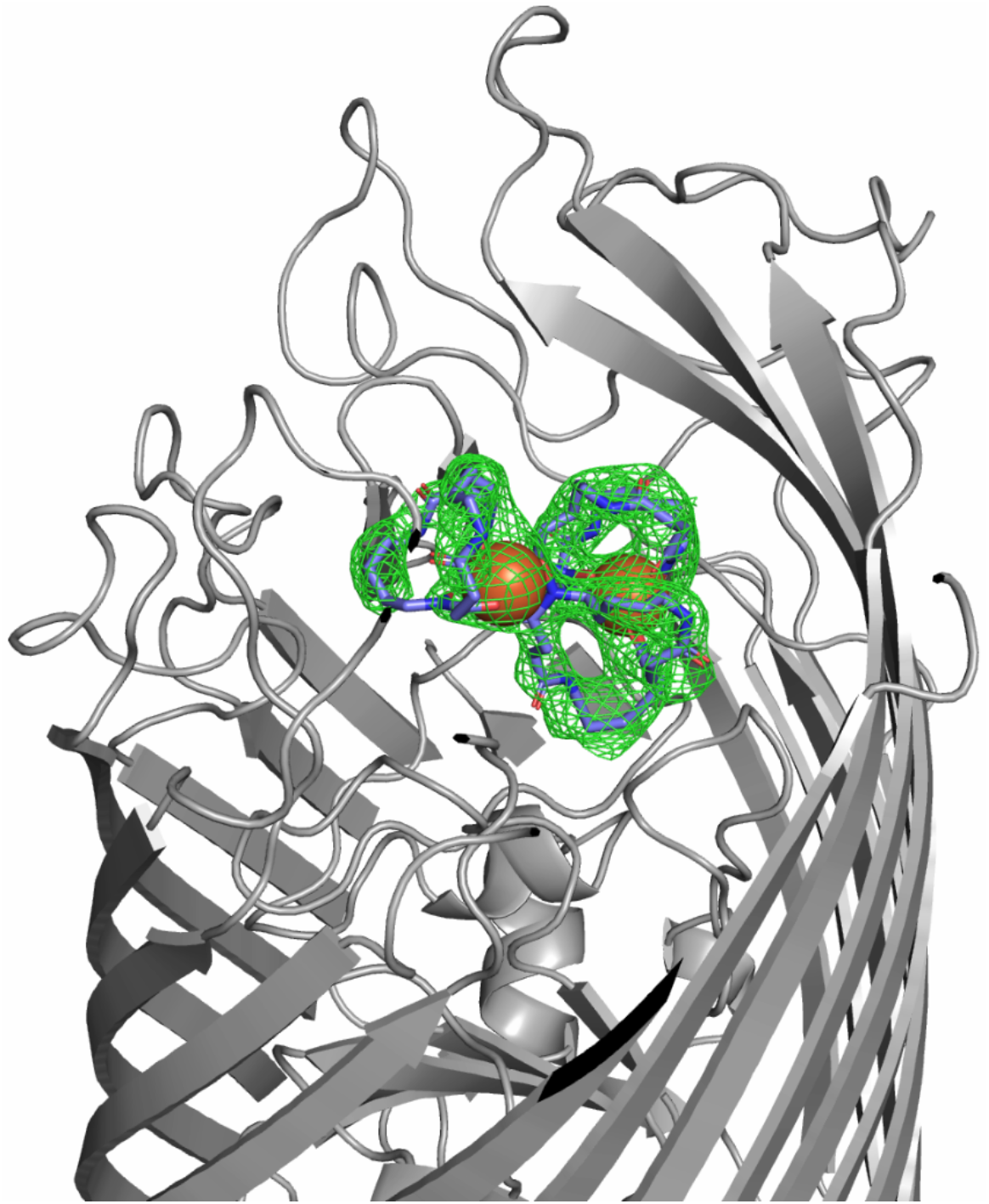
OMIT electron density map for bisucaberin_3_(blue)-Fe^3+^_2_ (orange)-FoxA (grey).

**Supplementary Data File 1:**
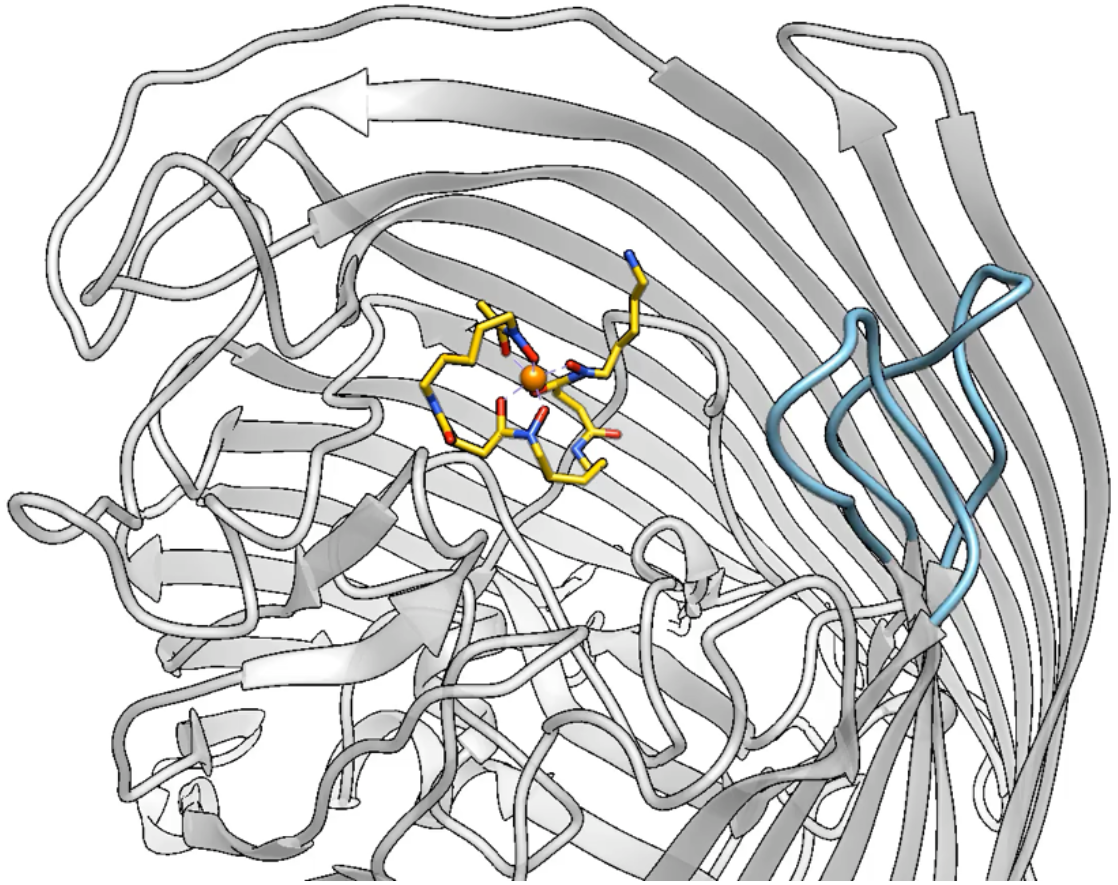
L7 and L8 fold inwards upon ferrioxamine B binding. FoxA is shown in grey. L7 and L8 are highlighted in blue whereas ferrioxamine is shown in yellow with heteroatoms in different colours. Conformational changes were modeled from PDB 6I96 and 6I97 and depicted using Chimera with the morph conformations tool.

